# A rapid, highly sensitive and open-access SARS-CoV-2 detection assay for laboratory and home testing

**DOI:** 10.1101/2020.06.23.166397

**Authors:** Max J. Kellner, James J. Ross, Jakob Schnabl, Marcus P.S. Dekens, Robert Heinen, Irina Grishkovskaya, Benedikt Bauer, Johannes Stadlmann, Luis Menéndez-Arias, Andrew D. Straw, Robert Fritsche-Polanz, Marianna Traugott, Tamara Seitz, Alexander Zoufaly, Manuela Födinger, Christoph Wenisch, Johannes Zuber, Vienna Covid-19 Detection Initiative (VCDI), Andrea Pauli, Julius Brennecke

## Abstract

Global efforts to combat the Covid-19 pandemic caused by SARS-CoV-2 still heavily rely on RT-qPCR-based diagnostic tests. However, their high cost, moderate throughput and reliance on sophisticated equipment limit widespread implementation. Loop-mediated isothermal amplification after reverse transcription (RT-LAMP) is an alternative detection method that has the potential to overcome these limitations. We present a rapid, robust, sensitive and versatile RT-LAMP based SARS-CoV-2 detection assay. Our forty-minute procedure bypasses a dedicated RNA isolation step, is insensitive to carry-over contamination, and uses a hydroxynaphthol blue (HNB)-based colorimetric readout, which allows robust SARS-CoV-2 detection from various sample types. Based on this assay, we have substantially increased sensitivity and scalability by a simple nucleic acid enrichment step (bead-LAMP), established a pipette-free version for home testing (HomeDip-LAMP), and developed open source enzymes that can be produced in any molecular biology setting. Our advanced, universally applicable RT-LAMP assay is a major step towards population-scale SARS-CoV-2 testing.

## Introduction

The Covid-19 pandemic poses unprecedented global health and economic challenges. The underlying Coronavirus Disease 2019 (Covid-19) is caused by infection with the single-stranded, positive-sense RNA beta-coronavirus SARS-CoV-2 (Gorbalenya et al., 2020). Without effective treatment, and with a large fraction of the world population not yet vaccinated, efforts to contain the spread of SARS-CoV-2 rely on systematic viral testing, contact tracing and isolation of infected individuals (Ferretti et al., 2020). Since SARS-CoV-2 carriers can be asymptomatic despite being infectious, a key challenge is to develop affordable and scalable technologies that enable population-wide testing (L. Zou et al., 2020). The gold-standard technique to detect an acute SARS-CoV-2 infection relies on nucleic acid diagnostics by RT-qPCR, which has been the method of choice due to its large dynamic range and high specificity (Corman et al., 2020). However, the need for specialized equipment and associated high cost make this technology unsuitable for population-scale testing, low resource settings and home testing. Moreover, slow turn-around times of several hours limit the applicability of RT-qPCR-based testing for situations where rapid screening is needed (CDC, 2020).

Isothermal nucleic acid amplification techniques, such as RPA (Recombinase-based Polymerase Amplification) (Piepenburg, Williams, Stemple, & Armes, 2006) or LAMP (Loop mediated isothermal amplification) (Notomi et al., 2000), have great potential to fill the technological gap required for large scale testing strategies as they enable rapid nucleic acid diagnostics with minimal equipment requirement (Niemz, Ferguson, & Boyle, 2011). Coupled to a reverse transcriptase step that converts viral RNA into single stranded DNA, several LAMP protocols for SARS-CoV-2 detection have been developed and applied to patient testing (Anahtar et al., 2020; Butler et al., 2020; Rabe & Cepko, 2020). Innovations such as a colorimetric read-out or the combination of RT-LAMP with specific CRISPR-Cas enzymatic detection has further simplified the assay and enhanced specificity, respectively (Zhang, Ren, et al., 2020; Broughton et al., 2020). However, several challenges remain, especially in terms of assay robustness, compatibility with crude patient samples, limitations in sensitivity, compatibility with home testing setups, and access to the patent-protected gold-standard RT-LAMP enzymes, which poses a central bottleneck for low-income countries.

Here, we present a highly versatile RT-LAMP assay that overcomes the afore mentioned limitations of current isothermal SARS-CoV-2 detection methods. We adopted an approach to greatly reduce the risk of carry-over contamination for SARS-CoV-2 testing, increased the robustness of the assay across all tested sample types and buffer conditions by using hydroxynaphthol blue (HNB) as colorimetric readout, boosted sensitivity by at least ten-fold by combining RT-LAMP with a simple RNA enrichment procedure, benchmarked a pipette-free method that enables sensitive and specific detection of SARS-CoV-2 in home settings, and finally present a powerful RT-LAMP assay that builds exclusively on open-source enzymes.

## Results

### A rapid, sensitive and specific RT-LAMP setup for SARS-CoV-2 detection

SARS-CoV-2 diagnostics relies on detection of viral RNA through reverse transcription and subsequent amplification of small parts of the 30 kilobase viral genome. Considering that in SARS-CoV-2 infected human cells the various subgenomic viral RNAs are expressed at different levels (Kim et al., 2020), we benchmarked six published SARS-CoV-2 specific primer sets (see Materials and Methods) based on their reported high sensitivities targeting different regions of the viral genome: the 5’-located ORF1ab gene, the envelope E gene and the most 3’-located N gene encoding the nucleocapsid protein (Figure 1A) (Broughton et al., 2020; Rabe & Cepko, 2020; Zhang, Odiwuor, et al., 2020; Zhang, Ren, et al., 2020). We used RNA extracted from nasopharyngeal swabs obtained from Covid-19 patients or confirmed SARS-CoV-2 negative individuals (negative controls) and a SARS-CoV-2 RNA standard to determine primer specificity and sensitivity in RT-LAMP reactions with fluorometric real-time readout. None of the six primer sets resulted in non-specific amplification within the first 50 minutes in negative controls. In contrast, when using patient RNA or synthetic SARS-CoV-2 standard as input, robust target amplification occurred after 10-20 minutes (Figure 1B, S1A, B). We conclude that RT-LAMP alone, without additional detection steps, is highly specific in complex human RNA samples. Throughout this study, we therefore recorded fluorescent real-time measurements or performed endpoint analyses after 30-35 minutes reaction time.

**Figure 1:**
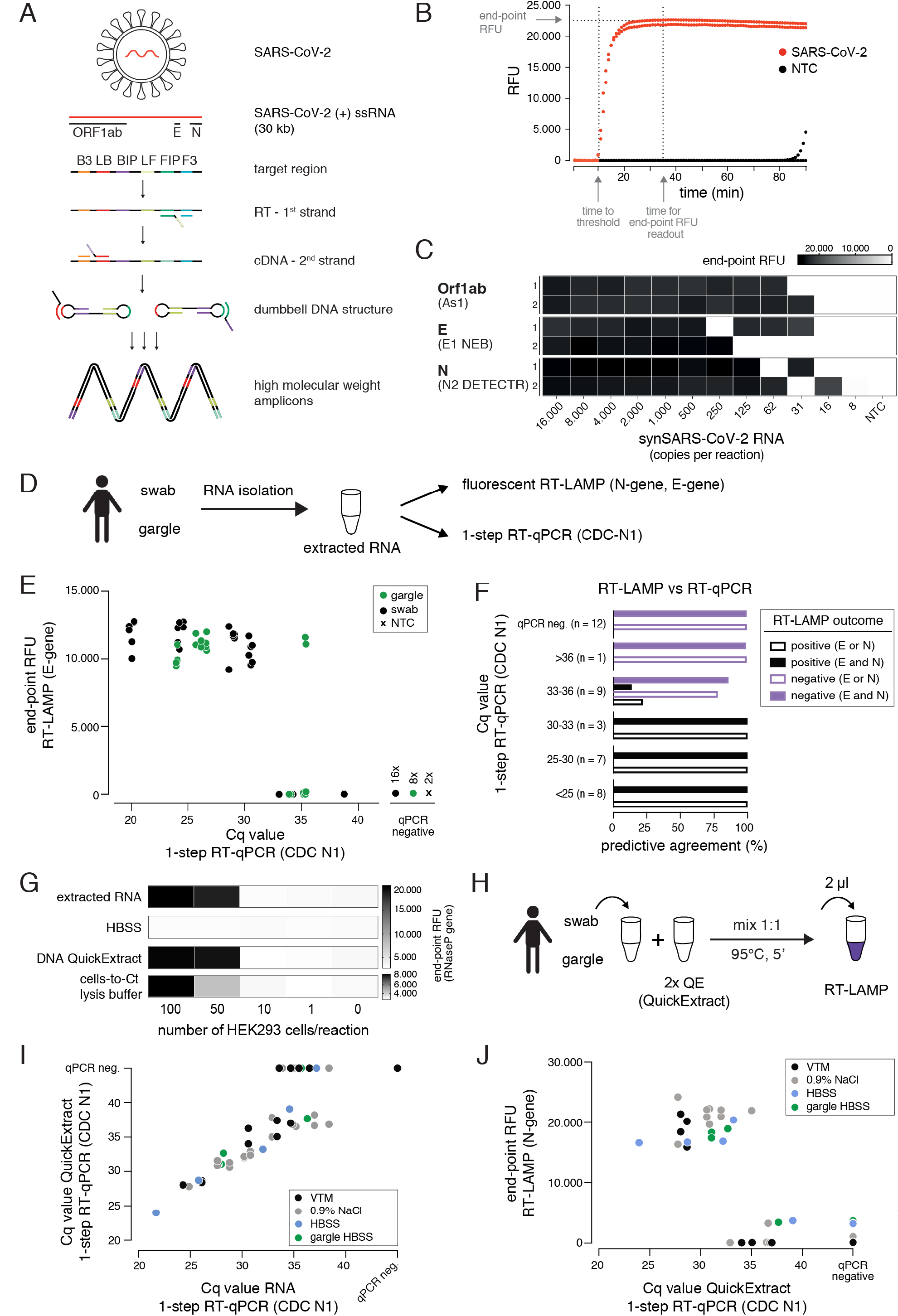
A sensitive, robust RT-LAMP assay compatible with crude patient samples. **A)** Schematic illustrating loop-mediated amplification (LAMP) of SARS-CoV-2 RNA and the regions targeted in this study (Orf1ab, E and N genes; depicted above). Each target region is recognized by a defined set of primers (B3, LB, BIP, LF, FIP, F3). The RNA template (red) is reverse transcribed and displaced after first-strand synthesis; the outer primer binding sites are added in the subsequent amplification step. The resulting dumbbell DNA structure acts as template for further rounds of amplification, ultimately leading to high molecular weight amplicons. **B)** Readout of a real-time fluorescence RT-LAMP reaction using 500 copies of synthetic SARS-CoV-2 (red) or water as non-targeting control (NTC, black) as input. ‘Time to threshold’ indicates the time at which the fluorescence value reaches threshold level (equivalent to Cq value in RT-qPCR assays), ‘end-point RFU’ indicates the fluorescence value (FAM filter set, absorption/emission at 494 nm/518 nm) after 35 minutes reaction time (used throughout this study unless indicated otherwise); RFU: relative fluorescence units. **C)** Performance of the three top primer sets for RT-LAMP-based SARS-CoV-2 detection. End-point relative fluorescence units (RFUs) of RT-LAMP reactions (in duplicates) using the indicated primer sets and serially diluted synthetic SARS-CoV-2 RNA standard as input. Water was used as no-target control (NTC). **D)** Cartoon indicating the workflow for SARS-CoV-2 detection by either RT-LAMP or 1-step RT-qPCR from patient samples (nasopharyngeal swab or gargle) with prior RNA isolation. **E)** Comparison of RT-LAMP and RT-qPCR performance. Plotted are RT-LAMP end-point fluorescence values after 35 minutes versus the respective RT-qPCR Cq values. RNA was derived from gargle (green) or nasopharyngeal swabs (black); two notarget controls were included (black cross). Reactions in which no amplification was recorded are labelled as qPCR negative. **F)** Predictive agreement between RT-LAMP and 1-step RT-qPCR assays. Shown are percentages of positive (detected in RT-LAMP and RT-qPCR, black bars) and negative (not detected in either RT-LAMP or RT-qPCR, purple bars) predictive agreement for sample groups (defined by RT-qPCR-derived Cq values) between RT-LAMP (using E- and/or N-gene primers) and 1-step RT-qPCR. **G)** Performance of different crude sample preparation methods in RT-LAMP. Shown are end-point relative fluorescence units (RFUs) for RT-LAMP reactions targeting human RNAseP on sample inputs derived from defined numbers of HEK293 cells mixed 1:1 with indicated 2x buffers (extracted RNsA served as a positive control). **H)** Cartoon indicating the workflow for RT-LAMP using QuickExtract crude lysate as sample input. **I)** Comparison of QuickExtract crude sample input versus extracted RNA as input using 1-step RT-qPCR. Covid-19 patient nasopharyngeal swabs or gargle samples (color coded according to the indicated collection medium) were either processed with the QuickExtract workflow (crude sample input) or RNA was extracted using an automated King Fisher RNA bead purification protocol. Reactions in which no amplification was recorded are labelled as qPCR negative. **J)** Performance of RT-LAMP with QuickExtract treated crude Covid-19 patient sample input (same samples as in I). Depicted is the correlation of Cq values from RT-qPCR performed on QuickExtract treated samples versus corresponding end-point relative fluorescence units (RFUs) from RT-LAMP reactions.

Three primer sets enabled SARS-CoV-2 detection down to ~16 copies per reaction (~8 copies/μl sample input): As1, E1 (NEB) and N2 (DETECTR) targeting the Orf1ab, E- and N-gene, respectively (Figure 1C, S1B) (Broughton et al., 2020; Rabe & Cepko, 2020; Zhang, Ren, et al., 2020). As previously reported (Anahtar et al., 2020; Rabe & Cepko, 2020; Zhang, Odiwuor, et al., 2020), RT-LAMP reactions with less than ~100 target molecules exhibited stochastic on-off outcomes. We therefore defined 100 copies per reaction as our robust limit of detection. When tested with a diluted patient sample, the E1 (NEB) and N2 (DETECTR) primers performed best with robust detection of SARS-CoV-2 up to Cq values of ~33 (sporadic detection up to Cq 35) (Figure S1B).

We next compared our RT-LAMP setup with one-step RT-qPCR, using RNA isolated from naso-pharyngeal swabs or gargle lavage from Covid-19 patients as input (Figure 1D-F). Using E1 or N2 primer sets for RT-LAMP, we achieved sensitive and specific detection of SARS-CoV-2 in patient samples with RT-qPCR measured Cq values of up to ~35 (~25 copies per reaction), independent of the patient sample type (Figure 1E). We obtained 100% positive predictive agreement rates between RT-LAMP and RT-qPCR up to Cq 33 (~100 copies per reaction) and 100% negative predictive agreement rates for qPCR negative samples (Figure 1F). E1 and N2 primer sets performed equally well, with a robust limit of detection of Cq 33-34 (Figure 1F). As shown recently (Zhang, Ren, et al., 2020), different primer sets can be combined in RT-LAMP reactions in order to reduce the false negative rate caused by suboptimal sample quality or by mutations in the viral genome coinciding with primer binding sites (Artesi et al., 2020).

A major bottleneck of certified RT-qPCR-based SARS-CoV-2 nucleic acid detection assays is their dependence on time-consuming and expensive RNA purification from patient samples. Inspired by recent findings (Ladha, Joung, Abudayyeh, Gootenberg, & Zhang, 2020; Rabe & Cepko, 2020), we assessed whether direct sample input/lysis conditions are compatible with sensitive RT-LAMP. Besides simple heat inactivation, we tested two previously published lysis and sample inactivation buffers, namely DNA QuickExtract (Lucigen) (Ladha et al., 2020) and the ‘Cells-to-Ct’ lysis buffer (Joung et al., 2017). To assess different lysis conditions, we compared crude lysates from serially diluted HEK293 cells to isolated RNA from equivalent numbers of cells as input for RT-LAMP reactions targeting the human reference gene RNaseP POP7 (Figure 1G). A five-minute incubation of patient samples with QuickExtract at 95°C performed equally well compared to a standard Trizol RNA extraction step (Ladha et al., 2020) (Figure 1G). Follow-up experiments substantiated that QuickExtract, in combination with heat treatment, preserves RNA integrity even under conditions where exogenous RNase A was added (Figure S2).

To benchmark QuickExtract solution on Covid-19 patient samples, we performed RT-qPCR on either purified patient RNA or crude QuickExtract lysate (Figure 1H). Irrespective of the sample type (swab or gargle), we observed a strong agreement between the corresponding RT-qPCR measurements (Figure 1I). Only samples with very low viral titers (high Cq values) became undetectable in the QuickExtract samples, presumably as ~20-fold less patient material equivalent was used compared to reactions using isolated RNA as input (Figure 1I). Importantly, RT-LAMP performed equally well to extracted RNA when using QuickExtract crude sample input across different transport media and different sample types (swabs in viral transport medium (VTM), swabs in 0.9% NaCl, swabs or gargle in HBSS buffer), with a limit of RT-qPCR-measured Cq values of 33 (~100 copies) and identical predictive performance rates (Figure 1J). No false positives were observed, demonstrating the high specificity and sensitivity of RT-LAMP on crude samples lysed and inactivated with QuickExtract solution. Heat inactivation with QuickExtract, in combination with fluorescent detection of the RT-LAMP reaction, is therefore a rapid method to detect SARS-CoV-2 in diverse patient samples.

### An efficient cross-contamination prevention system

LAMP results in the billion-fold amplification of target molecules. This poses a serious yet rarely mentioned risk, as only minor work-place or reagent contaminations with LAMP reactions will translate into large numbers of false positive assays (Kwok & Higuchi, 1989). Inspired by previous studies, we tested whether RT-LAMP based SARS-CoV-2 detection can be combined with a contamination prevention system that utilizes dUTP and thermolabile Uracil DNA Glycosylase (UDG) (Hsieh, Mage, Csordas, Eisenstein, & Tom Soh, 2014; Tang, Chen, & Diao, 2016). In this system, dUTP is incorporated into LAMP amplicons making them susceptible for Uracil-base cleavage in subsequent LAMP reactions containing the UDG enzyme (Figure 2A). To mimic carry-over contaminations from amplicons of prior LAMP reactions, we supplemented pre-RT-LAMP reactions (based on the key enzymes RTx and *Bst* 2.0) with dUTP, followed by dilution and addition to reactions in the presence versus absence of thermolabile UDG (Figure 2A). Thermolabile UDG is active at room temperature yet completely inactivated at temperatures above 50°C. In the absence of UDG, addition of a one billion-fold diluted pre-LAMP product resulted in indistinguishable signal in target vs. non-target conditions, illustrating the danger of cross-contamination. In contrast, in the presence of UDG, a 5-minute pre-incubation step at room temperature reduced the amplifiable carry-over product by more than 1,000-fold, enabling specific detection in the presence of considerable cross-over contamination product (Figure 2B). We conclude that the dUTP/UDG system is compatible with RT-LAMP reactions based on *Bst* 2.0 and RTx and profoundly lowers the risk of false positives.

**Figure 2:**
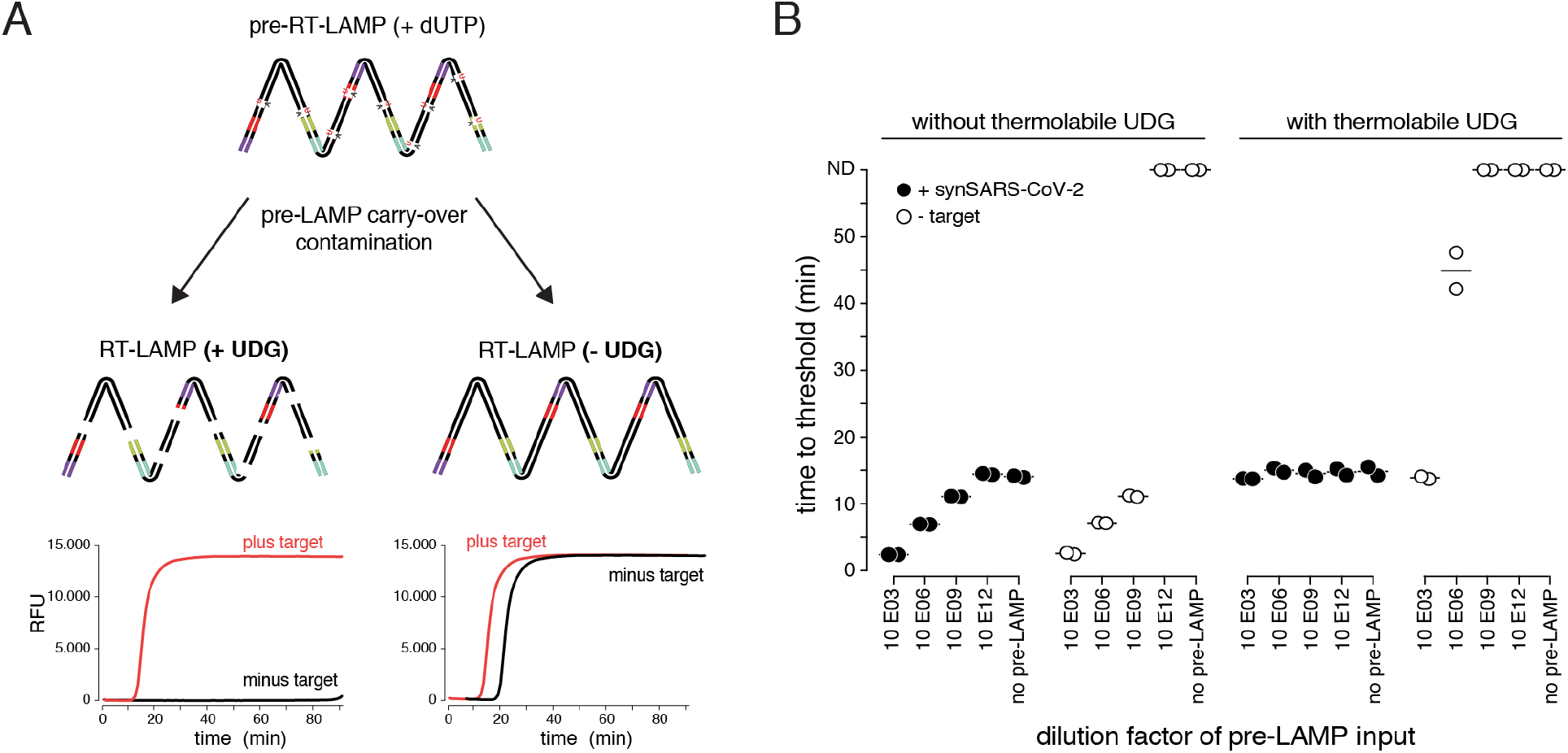
The dUTP/UDG system prevents carry-over cross-contamination in RT-LAMP. **A)** Schematic depicting the principle of the dUTP/UDG system in preventing carry-over contamination. dUTP is incorporated into LAMP amplicons in a primary reaction (pre-RT-LAMP). dUTP containing LAMP products carried over into a subsequent reaction (RT-LAMP) are cleaved by UDG prior to LAMP-based amplification, making them unavailable as amplification templates. This allows robust discrimination between target and no-target control (left), which is challenged by cross-over contamination in the absence of UDG-mediated cleavage (right). **B)** The dUTP/UDG system minimizes cross-over contamination. Shown are performances (time to threshold) of RT-LAMP reactions in the absence (left) or presence (right) of thermolabile UDG when using synthetic SARS-CoV-2 (filled circles) or water (open circles) as input. Reactions were supplemented with the indicated dilution of a dUTP-containing pre-LAMP reaction. All reactions were performed in duplicates.

### A robust colorimetric RT-LAMP readout compatible with various input samples

So far, we used real-time fluorescence (based on an intercalating DNA dye) to assess RT-LAMP-based target amplification. Given its dependency on specialized equipment, this detection method is prohibitive for low-resource settings or home-testing. Colorimetric detection resulting in a visual color change upon target DNA amplification provides an attractive, low-cost alternative (Goto, Honda, Ogura, Nomoto, & Hanaki, 2009; Tanner, Zhang, & Evans, 2015). Two colorimetric concepts are compatible with RT-LAMP: First, pH dependent dye indicators such as Phenol Red induce a color change from pink to yellow when the pH value of the reaction decreases upon DNA amplification (Tanner et al., 2015). Due to its pronounced color change, this is the most commonly used readout for RT-LAMP assays. However, the pH-change dependent readout requires a weakly buffered reaction solution, which poses a great challenge when using crude sample inputs with variable pH. A second colorimetric assay utilizes metal ion indicators such as hydroxynaphthol blue (HNB), which changes color from purple to blue upon a drop in free Mg2+ ions, which form a Mg-pyrophosphate precipitate upon DNA amplification (Figure 3A) (Goto et al., 2009).

**Figure 3:**
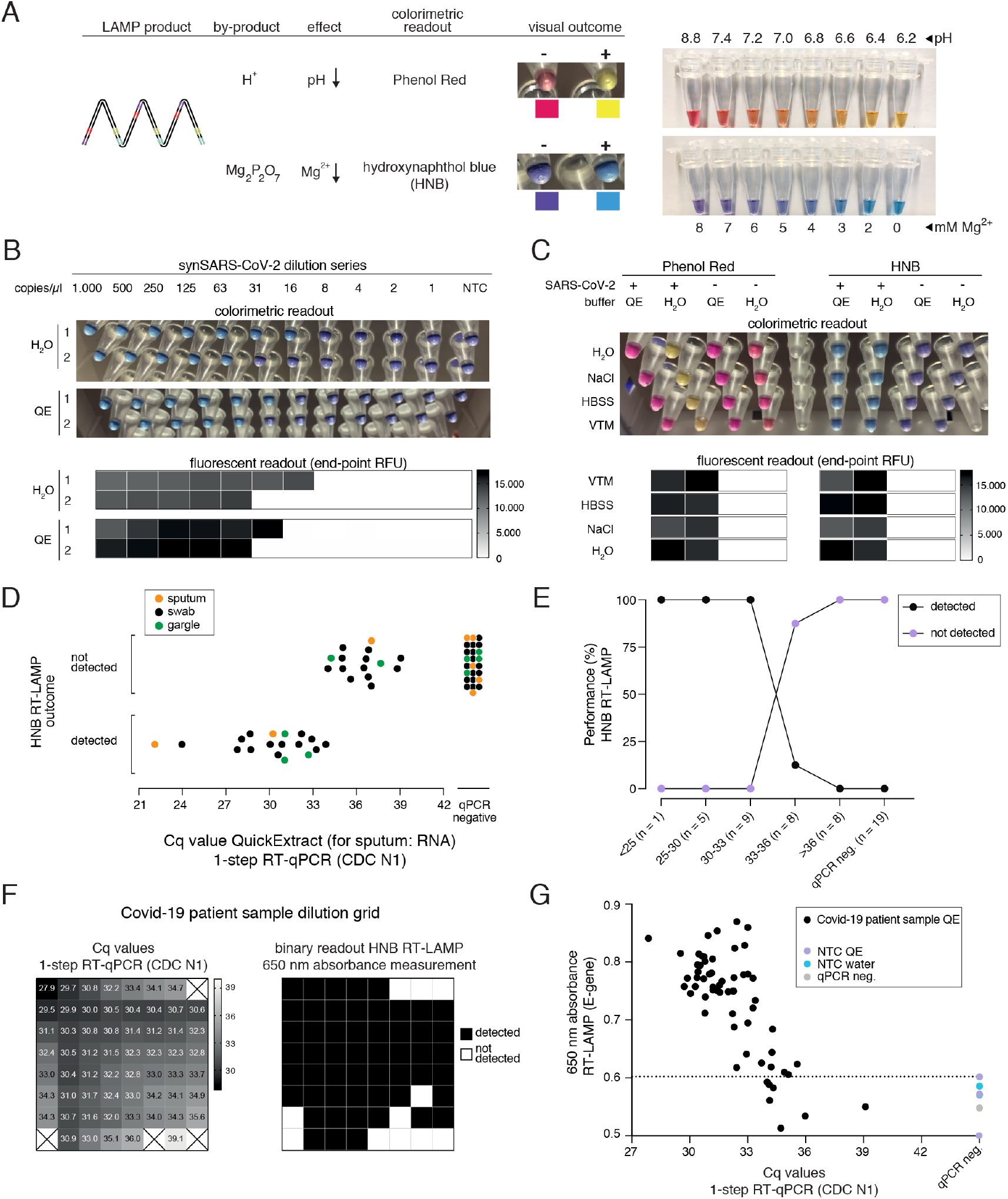
HNB RT-LAMP enables colorimetric SARS-CoV-2 detection from crude patient samples. **A)** Schematic illustrating the properties of pH-sensitive (Phenol Red, top) and Mg^2+^ concentration sensitive (hydroxynaphthol blue, HNB, bottom) colorimetric readouts for LAMP. Phenol Red interacts with protons (H+) generated during DNA amplification, which causes a color change from pink/red to yellow (right: the color range of the Phenol Red-containing colorimetric RT-LAMP mastermix (NEB) at relevant pH values). Magnesium pyrophosphate precipitate produced during DNA amplification lowers the free Mg^2+^ concentration, causing a color change of the HNB dye from purple to sky-blue (right: the color range of solutions with HNB at relevant Mg^2+^ concentrations). **B)** Influence of QuickExtract on HNB RT-LAMP performance. Shown is the colorimetric HNB readout of RT-LAMP reactions (after 35 minutes; in duplicates) using indicated copy numbers of SARS-CoV-2 RNA standard in water or QuickExtract and the corresponding co-measured end-point fluorescence values (heatmaps are shown below). **C)** QuickExtract lysis buffer is compatible with HNB colorimetric readout but incompatible with Phenol Red colorimetric readout of RT-LAMP reactions. Shown are RT-LAMP reaction outcomes (upper panel: colorimetric readout after 35 minutes, lower panel: fluorescent end-point values) when using 500 copies of synthetic SARS-CoV-2 RNA standard in indicated sample media diluted 1:1 with water or 2x QuickExtract solution as input. **D)** HNB RT-LAMP performance on Covid-19 patient samples lysed in QuickExtract solution. Shown is the binary colorimetric HNB readout of RT-LAMP reactions (N gene) using indicated patient samples (sputum (orange), swab (black), gargle (green)) plotted against the corresponding Cq values from RT-qPCR. **E)** Predictive agreement between HNB RT-LAMP and RT-qPCR assays using patient samples lysed in QuickExtract solution. Samples were grouped according to their RT-qPCR Cq values, and the percentage of detected (black) and not detected (purple) samples (based on HNB RT-LAMP) of the total number of samples per group was plotted. **F)** Schematic illustrating the serial dilution grid of a Covid-19 positive patient sample (Cq of 28) in QuickExtract. The heatmap (left) indicates Cq values determined by 1-step RT-qPCR (values above Cq 40 are indicated by black crosses). The grid (right) indicates the binary read-out (black: detected; white: not detected) of HNB RT-LAMP as measured by 650 nm absorbance). **G)** Scatter plot showing HNB RT-LAMP performance (measured by 650 nm absorbance) versus qPCR-determined Cq values on the serial dilution grid shown in F, including no-target controls (NTC) and a Covid-19-negative patient sample (qPCR negative). Horizontal dashed line indicates the maximum absorbance obtained for any negative control (y = 0.602).

Colorimetric readout, either via Phenol Red or the HNB dye, can be performed simultaneously with the fluorescent readout in the same RT-LAMP reactions. When using synthetic SARS-CoV-2 standard in water as input, both colorimetric readouts mirrored the fluorescent results (Figure 3B, S3). However, when using crude QuickExtract lysate as input, the pH-dependent readout failed or was inconclusive despite successful LAMP-mediated target amplification as evidenced by the fluorescent readout (Figure 3C, S3). In contrast, the HNB-dependent color change was not affected by QuickExtract solution, even when mixed with various sample buffers such as VTM, NaCl or HBSS (Figure 3C). We suspect that the QuickExtract solution is strongly buffered thereby preventing the required pH change that is typically generated during LAMP.

When tested in a clinical setting, RT-LAMP coupled to the HNB readout enabled robust detection of SARS-CoV-2 in patient samples with RT-qPCR values of up to ~34 (corresponding to ~50 copies per reaction of reference standard) with no false positives and 100% positive predictive agreement up to Cq 33 (~100 copies per reaction) (Figure 3D, E, S3A). The detection outcome was independent of the sample type, and we successfully used QuickExtract lysate from nasopharyngeal swabs, gargle solution or sputum samples (Figure 3D, S3B). We conclude that pH-independent dye formats, such as HNB, are superior in colorimetric RT-LAMP detection assays where strongly buffered or slightly acidic crude sample preparations are used as inputs.

To accurately determine the sensitivity threshold of HNB RT-LAMP, we generated a systematic dilution series of a positive Covid-19 patient sample in QuickExtract and used absorbance at 650 nm in a microplate reader to unambiguously determine the color change (Goto et al., 2009). We tested all dilutions by RT-qPCR and HNB RT-LAMP in parallel (Figure 3F, G). When considering samples with 650 nm absorbance values higher than for any co-measured negative control, HNB RT-LAMP allowed specific detection of samples up to Cq 34.9 with no false positives (Figure 3F, G). We conclude that, while read-out by fluorescence is the method of choice for high-throughput settings due to the higher dynamic range, direct absorbance measurement of the HNB-induced color change offers an attractive, alternative readout for large numbers of RT-LAMP reactions performed in parallel.

### An RT-LAMP assay with drastically increased sensitivity

SARS-CoV-2 RT-LAMP assays are roughly ten-fold less sensitive than conventional RT-qPCR assays. When using crude samples as input (i.e. eliminating the RNA concentration step by a dedicated extraction), sensitivity is further lowered, resulting in a substantial false negative rate. To increase the sensitivity of our RT-LAMP assay, we set out to establish a simple and rapid nucleic acid enrichment step. We used carboxylated magnetic beads to concentrate RNA from QuickExtract lysates on the bead surface in the presence of crowding agents and salt (Hawkins, O’Connor-Morin, Roy, & Santillan, 1994). We further reasoned that, instead of eluting RNA from the beads, adding the RT-LAMP mix directly to the dry beads should increase the number of viral RNA molecules per reaction by orders of magnitude, depending on the sample input volume (Figure 4A). We tested this approach, termed bead-LAMP, by using either bead-enriched or non-enriched synthetic SARS-CoV-2 RNA in HeLa cell QuickExtract lysate as RT-LAMP input. Indeed, bead-LAMP using 50 μl QuickExtract lysate as input displayed an at least ten-fold increase in sensitivity, corresponding to a robust limit of detection of ~5 copies per reaction (4/4 replicates; Cq value of 37-38) (Figure 4B-D). In three out of four replicates, bead-LAMP enabled detection of as few as 2 copies/μl, and in one replicate even a single copy/μl of sample input could be detected (Figure 4B, C). Overall, bead-LAMP drastically improved performance for samples with low viral titers that were non-detectable with regular RT-LAMP (Figure 4B, C, E) and reached RT-qPCR-like sensitivity (Figure 4E). The fluorescence readout of the bead-LAMP reaction exhibited overall lower values yet similar kinetics as regular RT-LAMP (Figure S6B), indicating that bead-LAMP is compatible with real-time kinetic analysis alongside colorimetric end-point detection (Figure 4B, C). After bead enrichment the recovery rates of synthetic SARS-CoV-2 RNA determined by RT-qPCR ranged from 68-98%, showing the high efficiency of the approach (Figure S4).

**Figure 4:**
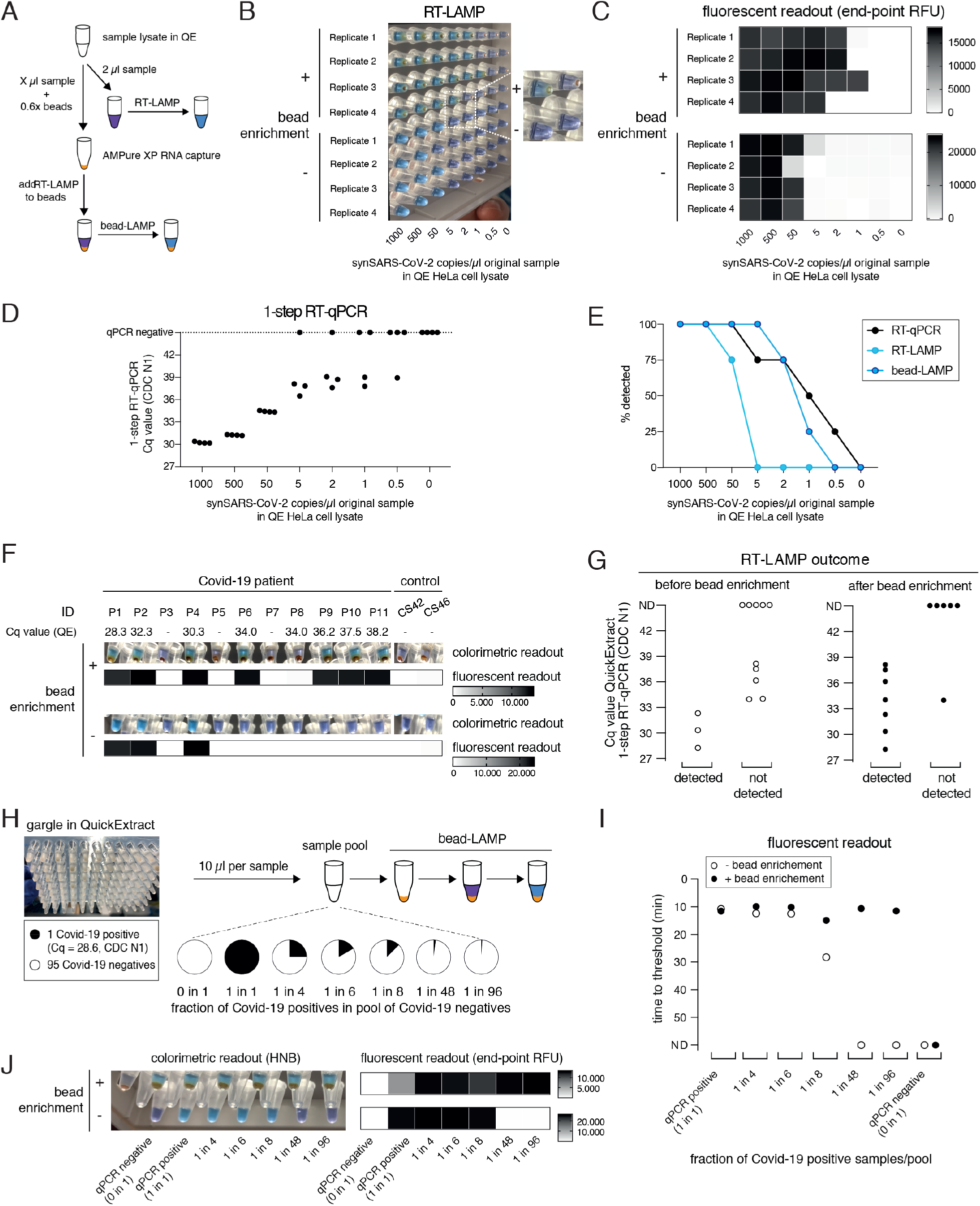
bead-LAMP increases sensitivity of RT-LAMP assays. **A)** Schematic illustrating the bead-LAMP workflow in comparison to the regular RT-LAMP workflow. AMPure XP RNA capture beads were used at 0.6x of the volume of the sample lysate (0.6x beads). **B)** Performance of bead-LAMP (+ bead enrichment) vs regular RT-LAMP (-bead enrichment) using a synthetic SARS-CoV-2 RNA standard spiked-in at the indicated concentration in the original sample into HeLa cell QuickExtract (QE) lysate. 50 *μ*l of crude sample in QE, adjusted to 100 *μ*l final volume with 1x HBSS was used for bead-enrichment. The image shows HNB end-point colorimetric readout of bead-LAMP and RT-LAMP reactions; a magnified view of four wells with beads (top: 5 copies/*μ*l is sky-blue; 2 copies/*μ*l is purple) and without beads (bottom: both are purple) is shown on the right. All reactions were performed in technical quadruplicates. **C)** End-point relative fluorescence units (RFUs), with or without prior bead enrichment for reactions shown in B. **D)** Performance of 1-step RT-qPCR using 2 *μ*l of the same crude sample preparations as used in B and C. **E)** Positive detection rates of 1-step RT-qPCR, RT-LAMP and bead-LAMP for reactions shown in B, C and D. **F)** Performance of bead-LAMP on a Covid-19 positive panel of patient samples in QuickExtract. The images depict the HNB colorimetric end-point readout, and the heatmaps underneath show co-measured end-point relative fluorescence units (RFUs) of RT-LAMP reactions, with or without prior bead enrichment, using eight Covid-19-positive and five negative samples as input (P1-P11, Covid-19 patient sample; CS42 and CS46, healthy controls). 100 *μ*l of crude sample in QE was used for bead-enrichment. Corresponding Cq values were obtained by measuring 2 *μ*l of the same QuickExtract (QE) patient samples by 1-step RT-qPCR prior to bead enrichment. **G)** Bead enrichment increases the sensitivity of RT-LAMP. Patient samples from F) were classified as detected or not detected based on the HNB RT-LAMP assay before (left, open circles) and after (right, filled circles) bead enrichment and plotted against their respective Cq values obtained from QuickExtract (QE) RT-qPCR (Cq values for qPCR negative samples are labelled as not detected, ND). **H)** Schematic illustrating the pooled testing strategy using bead-LAMP. A single Covid-19 positive patient gargle sample in QuickExtract (Cq ~28; black) was mixed with different amounts of 95 pooled SARS-CoV-2 negative samples (all in QuickExtract; white) yielding seven sample pools with indicated ratios of positive to negative samples. 40 to 100 *μ*l of crude sample in QE was used for bead-enrichment depending on the pool sizes. For lysate volumes smaller than 100 *μ*l, 1x HBSS was added to obtain a final volume of 100 *μ*l for bead-LAMP. **I)** Shown is the performance (measured as time to threshold) of bead-LAMP (filled circles) compared to regular RT-LAMP (open circles) on the patient pools defined in H. ND = not detected within 60 minutes of RT-LAMP incubation. **J)** Images showing the endpoint HNB colorimetric readout (left) and fluorescent readout (endpoint RFU; right) of samples measured in I) with or without prior bead enrichment.

We next tested bead-LAMP on a dilution series of a Covid-19 patient sample in QuickExtract (Cq of ~30) and observed a similar ten-fold increased sensitivity, corresponding to a limit of detection of ~Cq 37 in patient samples (Figure S5A). When performing bead-LAMP on individual Covid-19 patient samples, we found a dramatic improvement in the diagnostic performance. With the exception of one Covid-19 positive patient that we were not able to detect via RT-LAMP for unknown reasons, all qPCR positive samples (with Cq values up to ~38) were identified while no qPCR negative sample was detected (Figure 4F, G).

The boost in sensitivity opened the door for establishing a pooled RT-LAMP testing strategy. We mixed one crude Covid-19 positive patient gargle sample in QuickExtract (Cq ~28) with different volumes of a pool of 95 crude SARS-CoV-2 negative gargle samples in QuickExtract (Figure 4H, S5B). Each pool was tested by standard RT-LAMP and bead-LAMP. Without bead enrichment, pools with at least 12.5% (1 out of 8) of Covid-19 positive sample were identified. In contrast, bead-LAMP enabled detection of all pools containing SARS-CoV-2, even the pool containing just 1% (1 out of 96) of Covid-19 positive sample (Figure 4I, J, S5C). An independent experiment, in which we tested bead-LAMP on a dilution series of a Covid-19 positive patient of Cq ~30 in QuickExtract HeLa cell lysate, led to a similar conclusion: again, the pool containing only ~1% of the Covid-19 positive sample was detectable only with prior bead enrichment (Figure S5D-F). With merely 21 reactions (one entire 96-well plate pool, eight column pools, twelve row pools), a single positive patient of Cq ~30 can thus be detected amongst hundred individuals. We conclude that a cheap, fast (~5-10 minutes) and simple pre-enrichment step boosts the sensitivity of RT-LAMP ten-to fifty-fold, making this approach highly attractive for pooled testing strategies. Of note, the bead-based RNA enrichment step resulted in RT-LAMP reactions being fully compatible with the Phenol Red based colorimetric readout, even when QuickExtract lysates were used as input. While reactions without bead-enrichment failed to convert to yellow, the same input samples showed the characteristic color change when pre-purified via the bead-enrichment step (Figure S5G).

### A pipette-free RT-LAMP assay for home settings

The advancements presented so far provide a drastically improved SARS-CoV-2 detection assay regarding simplicity, robustness and sensitivity. However, our assay still required specialized laboratory equipment, such as precision pipettes or temperature-controlled incubators. We therefore explored approaches to adapt the HNB RT-LAMP protocol to home settings. In order to make RT-LAMP independent of precision pipettes, we adopted a previously reported strategy for sample clean-up and transfer using filter paper (Y. Zou et al., 2017). Using simple Whatman filter paper dipsticks (Figure 5A), we reliably detected SARS-CoV-2 RNA from Covid-19 patients with medium viral titers (Cq ~27) (Figure 5B). In addition, introducing a wash step with 130 mM sodium chloride solution increased the sensitivity and enabled SARS-CoV-2 detection in patient samples with Cq values ~32, mimicking the sensitivity of standard RT-LAMP assays (Figure 5B).

**Figure 5:**
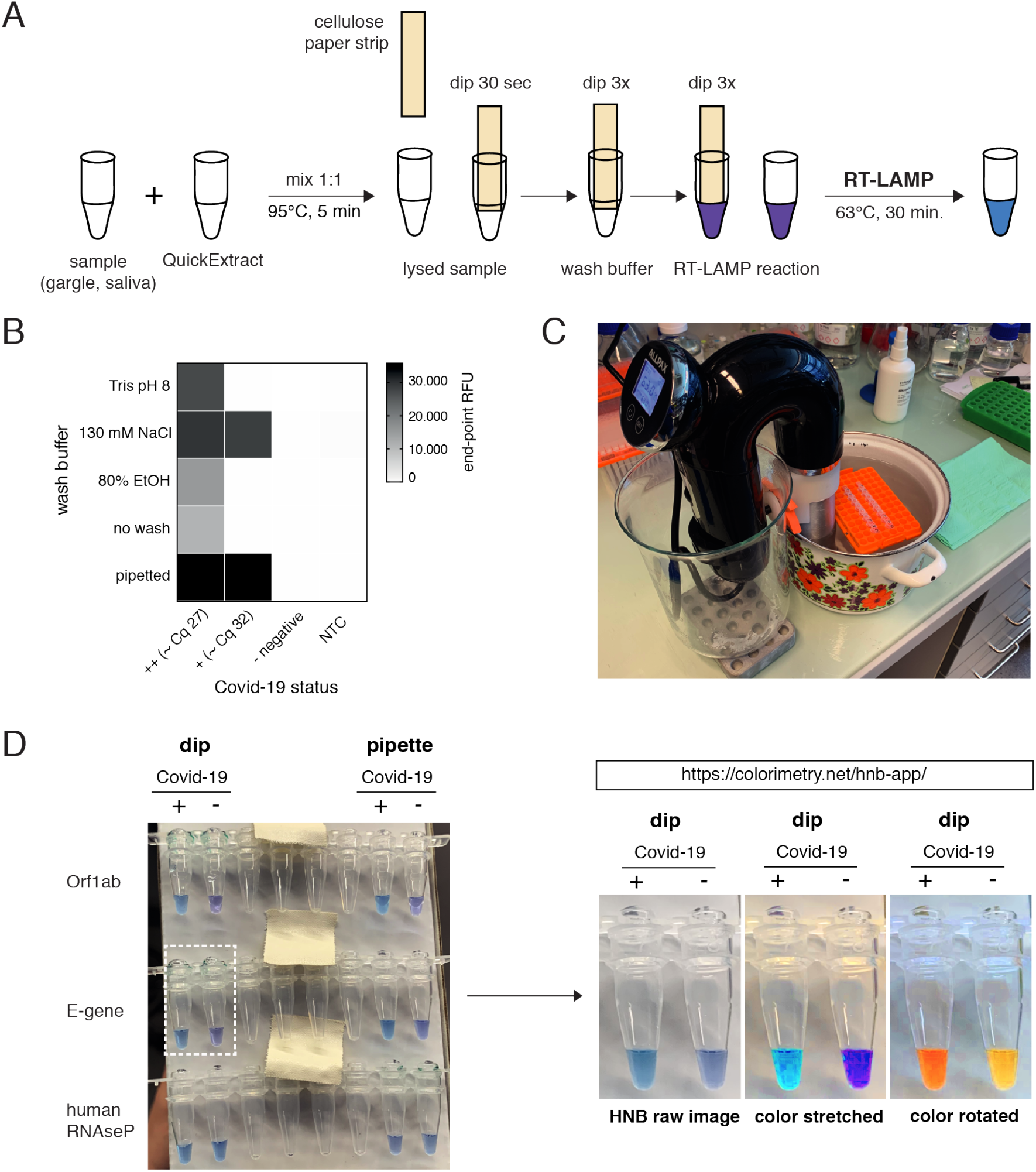
HomeDip-LAMP enables SARS-CoV-2 detection in low-resource and home settings. **A)** Schematic depicting the HomeDip-LAMP workflow. Samples are mixed 1:1 with QuickExtract lysis buffer and inactivated at 95°C for 5 minutes. Cellulose paper dipsticks are loaded by dipping into the crude sample for 30 seconds. After a brief washing step (3x dipping into wash buffer), RNA is released into pre-distributed RT-LAMP reaction mixes by 3x dipping. RT-LAMP reactions are performed in a water bath at 63°C and read out after 35 minutes. **B)** Influence of different wash conditions on SARS-CoV-2 detection using RT-LAMP with paper dipstick sample transfer. Heatmap showing end-point relative fluorescence units (RFUs) at 30 minutes of RT-LAMP reactions after transferring 2 *μ*l of high titer (++, Cq ~27), medium-to-low titer (+, Cq ~32) or negative Covid-19 patient samples in QuickExtract into 8 *μ*l of RT-LAMP reaction mix using cellulose paper dipsticks. Dipsticks were washed in between in indicated solutions or transferred without washing. A sample series where 2 *μ*l were transferred by pipetting (‘pipette’) is shown alongside (NTC = no target control). **C)** Image showing the water bath setup with a Sous Vide heater (black) for HomeDip-LAMP. Reaction tubes were kept upright and submerged using floating plastic pipette tip racks (orange). **D)** Detection of SARS-CoV-2 using HomeDip-LAMP. Left image shows true color readout (HNB dye) of HomeDip-LAMP (left 2 tubes) and pipetted LAMP (right 2 tubes) reactions using a Covid-19-positive (+) and -negative (-) patient sample in QuickExtract as input (35 minute end-point; water bath incubation at 63°C). Amplicons are indicated to the left; the human RNAseP amplicon served as positive control. The images to the right show color manipulations via the web-app (https://colorimetry.net/hnb-app/) of the two PCR tubes highlighted on the left for easier readout.

Due to their isothermal nature, RT-LAMP reactions require stable incubation temperatures of ~62-63°C. This can be provided using equipment ranging from high-end instruments to the most basal setup where boiling and room temperature water are mixed at a defined ratio and then kept insulated. We tested a commercially available sous-vide heater to create a temperature-controlled reaction environment (water bath) for home-based testing (Figure 5C). When combined with the filter paper-based sample clean-up and transfer method, this setup, termed HomeDip-LAMP, was able to accurately detect, within 35 minutes, two out of two viral genes in a Covid-19-positive patient gargle sample without false positives among Covid-19 negative gargle samples (Figure 5D). Detection accuracy and reaction speed matched HNB RT-LAMP reactions with pipetted sample input and laboratory equipment (Figure 5D). Moreover, a simple image manipulation (color stretch or rotation in HSL (Hue Saturation Lightness) space) via a custom-made web-app (https://colorimetry.net/hnb-app/) can convert the purple versus sky-blue color difference into more easily distinguishable outcomes, especially for untrained users (Figure 5D). Taken together, our findings provide a basis for the development of a simple SARS-CoV-2 detection platform, which can be implemented in any low-tech environment.

### A robust RT-LAMP assay using open source enzymes for SARS-CoV-2 detection

A critical bottleneck in population-scale testing efforts using RT-LAMP assays, especially in low-income countries, is the dependence on an existing and robust supply chain for the two enzymes, namely the reverse transcriptase (RT) and the *Bst* DNA Polymerase. All our assays so far relied on the patent-protected enzymes RTx, a thermostable RT, and *Bst* 2.0, an engineered variant of *Bst* LF, the Large Fragment of DNA polymerase I from *Geobacillus stearothermophilis* used in original LAMP assays (Notomi et al., 2000). Given that the sequences of neither enzyme are known, we set out to identify open-source enzymes that support RT-LAMP-based SARS-CoV-2 detection without compromising assay performance.

We first focused on the *Bst* DNA Polymerase and compared the engineered *Bst* 2.0 enzyme with the wildtype *Bst* LF counterpart. *Bst* LF exhibited similar overall reaction kinetics and sensitivity compared to *Bst* 2.0 (Figure 6A). Although reported to be more salt sensitive, *Bst* LF also allowed robust detection of SARS-CoV-2 from crude patient lysate in QuickExtract (Figure S6A). An important limitation of *Bst* LF is its incompatibility with the dUTP/UDG system due to its apparent inability to efficiently incorporate dUTP (Figure S6B). Nevertheless, considering the known protein sequence of wildtype *Bst* LF (GeneBank ID: AAB52611.1) and its open-source status, *Bst* LF is the enzyme of choice for settings where engineered *Bst* variants are not available or unaffordable (Bhadra, Riedel, Lakhotia, Tran, & Ellington, 2020; Phang, Teo, Lo, & Wong, 1995).

**Figure 6:**
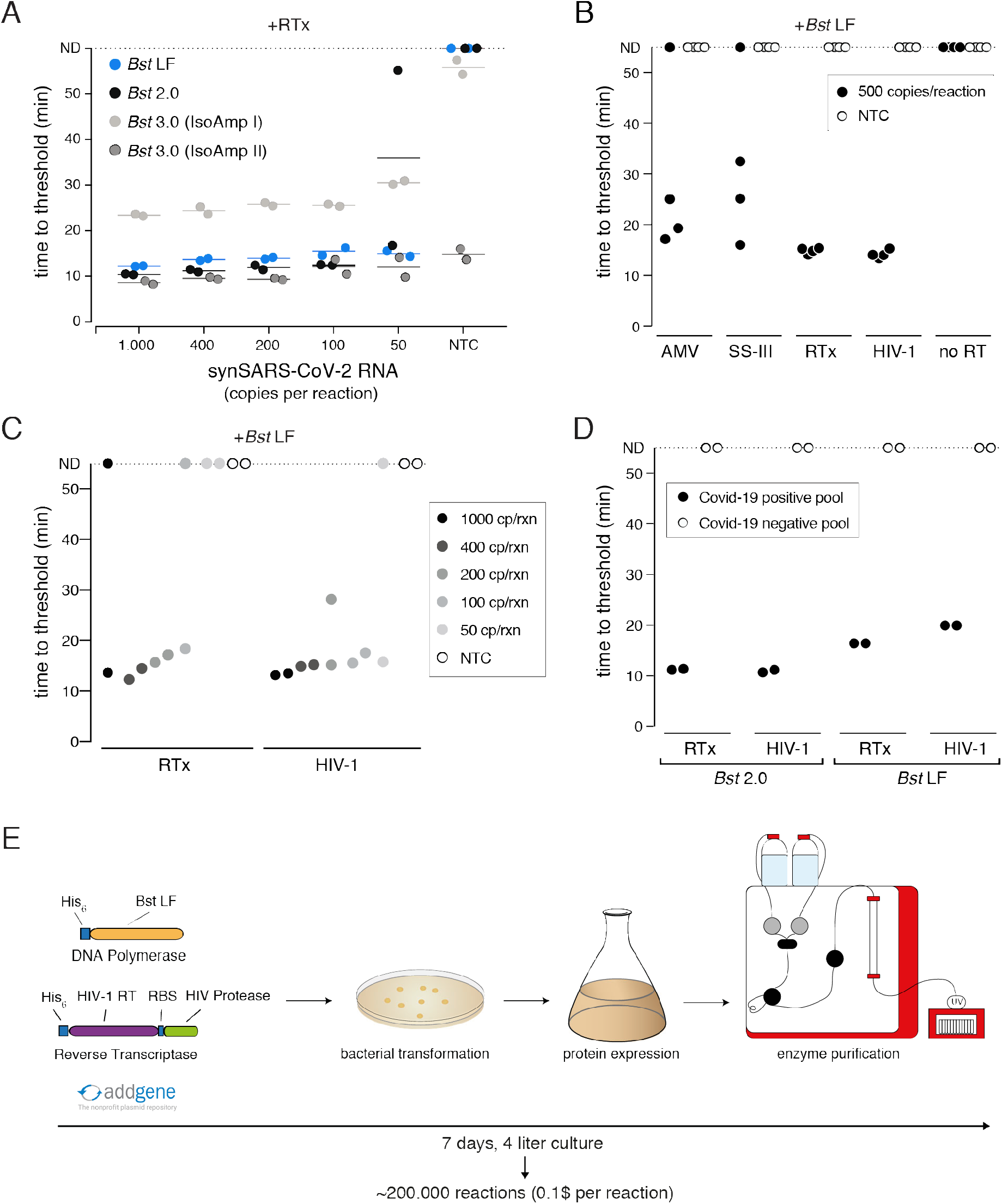
A sensitive RT-LAMP assay based on open-access enzymes. **A)** RT-LAMP performance (measured as ‘time to threshold’) of different *Bst* DNA polymerase variants in combination with NEB’s RTx reverse transcriptase on synthetic SARS-CoV-2 RNA standard. For reactions in which no amplification was recorded, ‘time to threshold’ is reported as ‘not detected’ (ND). Reactions were performed in duplicates; water was used as no-target control (NTC). **B)** RT-LAMP performance (measured as ‘time to threshold’) of different patent-protected (RTx, SuperScript III (SS-III)) and non-patent protected (AMV, HIV-1) reverse transcriptase enzymes in combination with *Bst* LF DNA polymerase on 500 copies/reaction of synthetic SARS-CoV-2 RNA standard. Reactions were performed in technical quadruplicates; water was used as no-target control (NTC). For reactions in which no amplification was recorded, ‘time to threshold’ is reported as ‘not detected’ (ND). **C)** RT-LAMP sensitivity performance (measured as ‘time to threshold’) of reactions containing NEB RTx or home-made HIV-1 RT, in combination with *Bst* LF DNA polymerase. Reactions contained different amounts of synthetic SARS-CoV-2 RNA standard. Reactions were performed in technical duplicates; water was used as no-target control (NTC). For reactions in which no amplification was recorded, ‘time to threshold’ is reported as ‘not detected’ (ND). **D)** Performance of RT-LAMP (measured as ‘time to threshold’) with different enzymatic compositions and QuickExtract patient sample as input. A pool of Covid-19 positive patient crude lysates (Pool N1-CDC, Cq 25) and a pool of Covid-19 negative crude lysates were tested in technical duplicates. For reactions in which no amplification was recorded, ‘time to threshold’ is reported as ‘not detected’ (ND). **E)** Deployment of open-access RT-LAMP. Bacterial expression plasmids can be obtained from Addgene (https://www.addgene.org/159148/, https://www.addgene.org/159149/). HIV-1 RT and Bst LF sufficient for ~220.000 reactions of RT-LAMP can be obtained within one week, starting from 4 liters of *E.coli* cultures.

We next examined the reverse transcription step that is required for the robust detection of RNA targets in RT-LAMP reactions. *Bst* polymerases are reported to exhibit intrinsic RT activity (Shi, Shen, Niu, & Ma, 2015). In particular, *Bst* 3.0 was engineered further from *Bst* 2.0 in order to display elevated intrinsic reverse transcriptase (RT) activity and increased amplification yield (patent US8993298B1). We thus tested *Bst* LF, *Bst* 2.0 and *Bst* 3.0 in LAMP reactions lacking the dedicated reverse transcriptase RTx. *Bst* LF showed no RT activity under our assay conditions (Figure S6C), and weak RT activity was observed for *Bst* 2.0 and *Bst* 3.0 when using universal Isothermal Amplification Buffer I (Figure S6C). In its optimized, higher-salt buffer (Isothermal Amplification Buffer II) *Bst* 3.0 yielded strong yet non-specific amplification irrespective of the presence of a dedicated RT (Figure 6A, S6C). A CRISPR-Cas12 collateral cleavage assay (Broughton et al., 2020) on the *Bst* 3.0 LAMP products revealed that *Bst* 3.0, in the absence of a dedicated RT enzyme, led to robust amplification of the synthetic standard down to 200 copies per reaction (Figure S6D-F). However, considering the patent-protection and the need for an additional detection step for amplicon-specific readout, we excluded *Bst* 3.0 as a possible entry point into an open-source RT-LAMP reaction. Instead, we concluded that a dedicated RT enzyme is required for efficient and specific target amplification.

To identify a thermostable, open-source RT that is active under the reaction conditions of LAMP, we first compared several RTs with the engineered RTx enzyme using synthetic SARS-CoV-2 RNA as template. We limited our test to RTs known to be active at elevated temperatures, namely AMV, Superscript III (SS-III) and a wildtype version of HIV-1 RT (Martín-Alonso, Frutos-Beltrán, & Menéndez-Arias, 2020). We found that wildtype HIV-1 RT worked equally well in terms of efficiency, speed and sensitivity as commercial RTx (Figure 6B, C), while AMV and SuperScript-III showed only limited RT activity (Figure 6B, S7A). Moreover, wildtype HIV-1 RT in combination with *Bst* LF was fully compatible with SARS-CoV-2 detection in crude patient samples (Figure 6D). As such, RT-LAMP with HIV-1 RT and *Bst* LF was able to detect SARS-CoV-2 RNA in a pool of Covid-19 positive patients but not in a pool of Covid-19 negative lysates (Figure 6D, S7B). Reaction speed in crude patient lysates was slightly reduced compared to the gold-standard RT-LAMP reaction using RTx and *Bst* 2.0, yet initiated still within 20 minutes (Figure 6D, S7B). We conclude that the combination of wildtype HIV-1 RT and *Bst* LF are fully able to perform RT-LAMP under our optimized reaction conditions with crude patient samples as input. These findings open the door for any laboratory to establish their own, home-made RT-LAMP reaction mix to enable SARS-CoV-2 and other pathogen testing.

## Discussion

RT-LAMP is an inexpensive and specific nucleic acid detection assay that provides test results in about thirty minutes. Its independence from specialized laboratory equipment and its compatibility with crude patient samples as well as colorimetric visual readout make it highly attractive for settings with limited resources or for population-scale testing. In this study, we systematically optimized every step of the RT-LAMP assay for crude sample input to make it more sensitive, more robust and simpler. Our improved assay holds the promise to contribute towards effective containment of the current SARS-CoV-2 pandemic.

Certified SARS-CoV-2 diagnostic workflows include an expensive and lengthy RNA isolation step. To circumvent this problem, several crude sample inactivation protocols have been developed that are compatible with direct downstream reverse transcription and amplification steps (Ladha et al., 2020; Myhrvold et al., 2018; Rabe & Cepko, 2020). While these advancements have simplified SARS-CoV-2 detection considerably, the required buffered solutions or reducing agents present a challenge for the commonly used pH indicator-based colorimetric LAMP readout. For example, strongly buffered lysis solutions such as QuickExtract are not compatible with Phenol Red dye detection, resulting in substantial false negative rates. Similarly, false positives have been reported for patient sample types with acidic pH such as saliva (Lalli et al., 2020). We employed the known metal indicator hydroxynaphthol blue (HNB) as a robust alternative for colorimetric detection of SARS-CoV-2 with no false positives and detection rates identical to highly sensitive fluorescent LAMP assays for any tested sample buffer (Figure 3, S3). HNB RT-LAMP, particularly when employed in combination with the dUTP/UDG cross-contamination prevention system (Figure 2), is therefore a highly robust and stream-lined assay suited for SARS-CoV-2 testing in home settings.

A major drawback of using patient samples directly for nucleic acid detection is the resulting drop in sensitivity. While an upstream RNA isolation step allows the concentration of viral template molecules, this is not the case for crude extraction methods. With a robust limit of detection of ~50-100 copies per reaction, RT-LAMP with crude patient sample input can only detect medium to high viral titers. Our development of bead-LAMP, a simple RNA enrichment protocol with magnetic beads, sets the stage for highly sensitive SARS-CoV-2 detection in samples from individual patients or patient pools (Figure 4). While similar to a recently reported protocol based on silica particles (Rabe & Cepko, 2020), our approach requires only a magnet and adds just 5-10 minutes to the standard protocol. Bead-LAMP does not require centrifugation and can be performed manually with a simple magnet, an automated magnetic particle processor like the KingFisher or on fully automated liquid handling platforms. It is especially suited for mass-scale pathogen surveillance via sample pooling strategies. Combined with the HNB colorimetric read-out, bead-LAMP allows for screening hundreds of individuals in pooled reactions in simple PCR strips. Bead-LAMP is also an attractive alternative to ultra-sensitive RT-qPCR when used on single patient samples (Figure 4F, G).

To illustrate the practical implications of assay sensitivity, we used our RT-qPCR data from nasopharyngeal swabs to calculate viral copies per entire clinical sample as done in (Wölfel et al., 2020). Figure S8A shows that the viral titer per sample ranges from ~650 to 2×10^8^ viral RNA copies. Grouping patient samples according to their ability to infect cells in culture (>10^6^ viral copies per swab) (Wölfel et al., 2020) further allows to separate them roughly into infectious and non-infectious groups. RT-qPCR coupled to RNA isolation is able to detect this enormous range of viral titers (down to 5 copies per reaction) (Corman et al., 2020). Without distinguishing between infectious or non-infectious individuals, RT-LAMP on purified RNA (limit of detection: 100 copies per RT-LAMP reaction) would detect ~ 80% of all infected individuals (Figure S8B). RT-LAMP on QuickExtract lysate directly prepared from crude sample lowers the number of detectable individuals to 64% or 50% when swab volumes of 0.5 ml or 3 ml are used, respectively. In contrast, the theoretical detection rate of bead-LAMP is ~92% relative to RT-qPCR on extracted RNA, when considering 100% RNA recovery on beads, and up to 86% when taking the average 77% recovery measured with our protocol (Figure S4) into account. Of note, bead-LAMP is more sensitive than RT-qPCR performed on crude QuickExtract lysate, which detects ~80% of infected individuals in our experiments. It is important to note that infectious patients are identified at 100% detection rate in all RT-LAMP detection formats, highlighting the relevance of RT-LAMP for population screening. Taken together, in combination with minimizing sample volumes of nasopharyngeal swabs, bead-LAMP, without a dedicated RNA purification step, reaches the sensitivity that is currently defined by gold-standard certified RT-qPCR assays. At the same time, bead-LAMP outperforms RT-qPCR in terms of speed, cost (~1.5 USD for commercial QuickExtract & RT-LAMP reagents plus 0.1 – 0.25 USD for commercial magnetic beads per reaction), and low equipment needs.

While bead-LAMP enables pooled testing, reliable and sensitive home-tests provide important alternative strategies in combating the Covid-19 pandemic (Taipale, Romer, & Linnarsson, 2020). Towards this end, we present a simple strategy for sample RNA binding and transfer using cellulose paper strips. With HomeDip-LAMP, SARS-CoV-2 detection can be performed in home settings without the use of pipettes (Figure 5). Only sample inactivation buffer, paper-strips, wash and reaction solution together with a stable heat-source such as a water bath are required. We envision that a combination of bead-LAMP with HomeDip-LAMP could be adapted for sensitive home testing. In such a combined approach, beads could be added to the inactivated sample, followed by binding to a magnetic rod and dipping as described for cellulose paper strips.

Establishing RT-LAMP based SARS-CoV-2 testing in developing countries is severely hampered by unreliable or non-existing supply chains. The gold-standard RT-LAMP enzymes *Bst* 2.0 and RTx are engineered and proprietary enzymes, making their on-site production impossible. In our tests, the wildtype *Bst* LF enzyme performed equally well to *Bst* 2.0, with the exception of dUTP incorporation. One of our most significant findings was the identification of a wildtype HIV-1 RT enzyme as an eye-level alternative to the RTx enzyme. *Bst* LF and HIV-1 RT can be recombinantly produced at high yields using simple molecular biology equipment (Boretto et al., 2001; Matamoros et al., 2005). The implications of our identification of open-access enzymes that support rapid and sensitive RT-LAMP are profound: in principle, any molecular biology lab in the world will be able to generate a robust and sensitive in-house RT-LAMP reaction mix.

In summary, our improvements over existing RT-LAMP workflows enable the robust, in-expensive and ultra-sensitive detection of SARS-CoV-2. Our findings provide the basis for future clinical performance studies with the ultimate goal to make ‘testing for everyone’ a reality. Combating the Covid-19 pandemic will require access to diagnostic tests in all countries (“The COVID-19 testing debacle,” 2020). Our establishment of an RT-LAMP assay using only open-access enzymes is a major step forward to meet precisely this need.

## Acknowledgements

This work would not have been possible without the enthusiastic support of the IMBA and IMP research institutes, as well as the many volunteers and partners of the VCDI, who came together to help and collaborate under the exceptional circumstances of the Covid-19 pandemic. We thank the Pauli and Brennecke groups as well as Peter Duchek and Arabella Meixner for bearing with us and sharing lab space, and M. Voichek, V. Deneke and D. Handler for support and discussions. We thank Nathan Tanner (NEB) for valuable discussions, sharing important information on LAMP technology and feedback on the manuscript, and Stuart Le Grice and Jennifer Miller (NIH/NCI) for helpful advice and reagents for expression of HIV-RTs. We are grateful to the Covid Testing Scaleup SLACK channel for openly sharing and exchanging information and the Covid-19 diagnostics team in Feng Zhang’s lab for discussions.

## Author contributions

MJK, JJR and JS designed, performed and analyzed all experiments; MPSD, RFP and RH processed patient samples; IG, BB, JS and LMA performed protein expression studies; ADS designed and programmed the colorimetry web-application; MT, TS, AZ, MF and CW provided patient samples; JZ, MF and CW coordinated clinical validation studies; JZ acquired project funding; the VCDI enabled and supported Covid-19 testing initiatives at the Vienna BioCenter; MJK, AP and JB conceived the project; AP and JB coordinated and supervised the project; MJK, AP and JB, with the help of JJR and JS, wrote the paper with input from all authors.

## Materials and Methods

### Clinical sample collection

Nasopharyngeal swabs were collected in 1.5-3 ml VTM, 0.9% NaCl solution or 1x HBSS (Gibco: 140 mM NaCl, 5 mM KCl, 1 mM CaCl_2_, 0.4 mM MgSO_4_-7H2O, 0.5 mM MgCl_2_-6H2O, 0.3 mM Na_2_HPO_4_-2H2O, 0.4 mM KH_2_PO_4_, 6 mM D-Glucose, 4 mM NaHCO_3_). Gargle samples were collected from swab-matched patients by letting individuals gargle for 1 minute with 10 ml of HBSS or 0.9% Saline solution. Sputum samples were prepared by mixing sputum material 1:1 with 2x Sputolysin solution (6.5 mM DTT in HBSS) and incubation at room temperature for 15 minutes. The study was approved by the Ethics Committee of the city of Vienna (EK-20-208-0920). Written consent was obtained from all patients.

### RNA extraction from patient material

Total RNA was isolated from 100 *μ*l of nasopharyngeal swabs or cell-enriched gargling solution using a lysis step based on guanidine thiocyanate (adapted from Boom et al. 1990) and 20 *μ*l of carboxylated magnetic beads (GE Healthcare, CAT: 65152105050450) applied in 400 *μ*l of Ethanol on the magnetic particle processor KingFisher (Thermo). After a 5-minute incubation at room temperature, DNA was digested with DNaseI for 15 mins at 37°C, followed by a series of wash steps. RNA was eluted from the beads in 50 *μ*l RNase free H2O for 5 minutes at 60°C.

### Crude sample inactivation using QuickExtract DNA solution

50 *μ*l of nasopharyngeal swabs, gargle solution or sputum sample were mixed 1:1 with 2x QuickExtract DNA extraction solution (Lucigen) and heat inactivated for 5 minutes at 95°C. Samples were then stored on ice until further use or frozen at −80°C.

### RT-qPCR

For detecting the viral N-gene via RT-qPCR, 1-step RT-qPCR was performed using the SuperScript III Platinum One-Step qRT-PCR Kit (Thermofisher) or Luna Universal One-Step RT-qPCR Kit (NEB) and 1.5 *μ*l of reference primer/probe sets CDC-N1 (IDT 10006713) or CDC-N2 (IDT 10006713) per 20 *μ*l reaction. Reactions were run at 55°C for 15 minutes, 95°C for 2 minutes, followed by 45 cycles of 95°C for 10 seconds and 55°C for 45 seconds in a BioRad CFX qPCR cycler. Each RT-qPCR reaction contained either 5 *μ*l (N-gene, extracted RNA) or 2 *μ*l (N-gene, QuickExtract lysate) of sample input per 20 *μ*l reaction.

### Fluorescent RT-LAMP

Fluorescent RT-LAMP reactions were set up using the NEB Warmstart RT-LAMP kit or individual enzymes. For reactions using the RT-LAMP kit, Warmstart RT-LAMP master mix (1x final, 2x stock) was mixed with primer solution (1x final, 10x stock) containing all six LAMP primers (B3, F3, LB, LB, FIP, BIP), LAMP dye (1x final, 50x stock) or Syto9 (1 *μ*M final, 50 *μ*M stock), sample and nuclease-free water. Primers were used at final concentrations of 0.2 *μ*M for F3/B3, 0.4 *μ*M for LB/LF (except for N2 DETECTR, LB/LF 0.8 *μ*M) and 1.6 *μ*M FIP and BIP. Typical final reaction volumes were 10 *μ*l or 20 *μ*l containing 2 *μ*l of sample.

For LAMP reactions using individual polymerases, RT-LAMP reactions were set up using NEB 1x Isothermal Amplification Buffer (*Bst* LF, *Bst* 2.0, *Bst* 3.0) or NEB 1x Isothermal Amplification Buffer II (*Bst* 3.0), 6 mM MgSO4 (8 mM final; 2 mM MgSO_4_ are present in Isothermal Buffer I), 0.3 U/*μ*l NEB Warmstart RTx, 0.32 U/*μ*l NEB *Bst* DNA polymerase (LF, 2.0 or 3.0), 1.4 mM of each dNTP (Larova, 25 mM of each dNTP stock solution), 1x fluorescent dye or 1 *μ*M Syto9, sample and nuclease-free water.

For LAMP reactions testing individual RT-enzymes, RT-LAMP reactions were set up using NEB 1x Isothermal Amplification Buffer (*Bst* LF, *Bst* 2.0, *Bst* 3.0) or NEB 1x Isothermal Amplification Buffer II (*Bst* 3.0), 6 mM MgSO4 (8 mM final; 2 mM MgSO_4_ are present in Isothermal Buffer I), 1.4 mM of each dNTP (Larova, 25 mM of each dNTP stock solution), 0.32 U/*μ*l NEB *Bst* DNA polymerase (LF, 2.0 or 3.0), 0.3 U/*μ*l, Warmstart RTx (NEB), 0.2 U/*μ*l AMV RT (NEB), 4 U/*μ*l SuperScript III (Thermofisher), 50 nM of home-made HIV-1 RT (BH10) diluted in 1x dilution buffer (TrisHCl pH 7.5, 50 mM NaCl, 0.5 mM TCEP) and 1x fluorescent dye or 1 *μ*M Syto9, sample and nuclease-free water.

Reactions were run at 63°C (62°C for N2 DETECTR and primer comparison) for 30-60 minutes in a BioRad CFX Connect qPCR cycler with SYBR readings every minute.

### Direct sample lysis buffer test

HEK293 cells were trypsinized and counted to make the appropriate dilutions in HBSS. The dilutions were mixed 1:1 with respective lysis buffers and treated as follows: Cells for no extraction were incubated for 5 min at room temperature. QuickExtract samples were incubated at 95°C for 5 min. Cells lysed in the home-made buffer (19.2 mM Tris–HCl (pH 7.8), 1 mM MgCl_2_, 0.88 mM CaCl_2_, 20 *μ*M DTT, 2% (wt/vol) Triton X-100) were incubated for 5 min at room temperature before incubation at 95°C for 5 min. For extracted RNA, RNA was purified from 1e5 HEK293 cells using standard Trizol RNA extraction and diluted to cell/reaction equivalents.

### dUTP/UDG contamination prevention system

Reactions were set up to contain NEB 1x Isothermal Amplification Buffer, 1.4 mM of each dATP, dCTP, dGTP, 0.7 mM dUTP, 0.7 mM dTTP, 6 mM MgSO4 (100 mM stock, NEB), 0.32 U/*μ*l NEB *Bst* 2.0 polymerase, 0.3 U/*μ*l NEB Warmstart RTx Reverse Transcriptase, 0.2 U/*μ*l NEB Antarctic thermolabile UDG, sample and nuclease-free water. Reactions were set up on ice and incubated at room temperature for 5 minutes before being transferred to 63°C to start RT-LAMP reactions under standard conditions described above. For demonstrating carry-over contamination, reactions either contained UDG (+UDG) or water (-UDG) and different amounts of pre-amplified RT-LAMP product (pre-RT-LAMP). Pre-RT-LAMP reactions were performed with dUTP, E-gene primer and 500 copies of Twist synthetic RNA standard for 60 minutes at 63°C. Serial dilutions were made by mixing 1 *μ*l of dUTP-containing pre-RT-LAMP product with 999 *μ*l of nuclease-free water to get 1e3-, 1e6-, 1e9- and 1e12-fold dilutions of pre-RT-LAMP, followed by addition of 2 *μ*l diluted pre-RT-LAMP product to dUTP/UDG RT-LAMP reactions.

### Colorimetric LAMP

For HNB colorimetric RT-LAMP detection, reactions were set up as in fluorescent RT-LAMP with the addition of 120 *μ*M HNB dye solution (20 mM stock in nuclease-free water). Phenol Red colorimetric reactions were performed using the NEB WarmStart colorimetric LAMP 2x master mix and the same final primer concentrations as in fluorescent RT-LAMP reactions. HNB and Phenol colorimetric reactions further contained 1x fluorescent LAMP dye (50x stock from LAMP kit) or 1 *μ*M Syto9 dye (50 *μ*M Stock) to measure fluorescence in parallel.

### Bead-LAMP

For bead enrichment, variable volumes of sample in QuickExtract (40 *μ*l up to 100 *μ*l) were adjusted to a final volume of 100 *μ*l with HBSS, mixed with 0.6x of beads (1:5 dilution of Agencourt RNAClean XP in 2.5 M NaCl, 10 mM Tris-HCl pH 8.0, 20% (w/v) PEG 8000, 0.05% Tween 20, 5 mM NaN_3_) and incubated for 5 minutes at room temperature. Beads were captured with a magnet for 5 minutes and then washed twice with 85% ethanol for 30 seconds. The beads were air dried for 5 minutes and then eluted directly in 20 *μ*l colorimetric HNB LAMP reaction mix containing 1x NEB WarmStart LAMP kit, 1x Fluorescent LAMP dye, 120 *μ*M HNB dye solution and 1x primer mix. No additional volume for dry beads was factored into the RT-LAMP reaction mix such that reactions were completed with nuclease free water to have final reaction volumes of 20 *μ*l.

As sample input for pooled bead-LAMP (Figure 4H-J, S5B-D), sample pools were prepared by mixing 10 *μ*l of a Covid-19 positive patient gargle sample in QuickExtract with different amounts of a Covid-19 negative gargle sample pool (n=95) in QuickExtract (10 *μ*l per sample). For pool volumes <100 *μ*l, the volume was filled up to 100 *μ*l with HBSS:QuickExtract (1:1); for pool volumes >100 *μ*l, an aliquot of 100 *μ*l were taken out after pooling for subsequent RT-LAMP or bead-LAMP. 40 *μ*l (matching the smallest pooled sample volume) of a Covid-19 positive or negative patient gargle sample were used as positive (qPCR positive) or negative (qPCR negative) controls, and also filled up to 100 *μ*l with HBSS:QuickExtract (1:1) before LAMP.

As sample input for the proof-of-concept experiment shown in Figure S5D-F, sample pools containing different numbers of Covid-19 positive patient gargle sample in QuickExtract were mixed with HeLa cell lysate in QuickExtract. The HeLa cell lysate was prepared by adding 500 *μ*l of HBSS and 500 *μ*l of 2x QuickExtract solution to a HeLa cell pellet containing one million cells, followed by cell lysis for 5 minutes at 95°C. The stock lysate of 1000 cells/*μ*l was then diluted in 1x heat inactivated QuickExtract buffer (diluted to 1x in HBSS) to a final concentration of 20 cells/*μ*l. This concentration was chosen as QuickExtract lysate from gargle or swabs roughly yields 200 pg/*μ*l of RNA or 20 cells/*μ*l. This Covid-19 negative QuickExtract lysate was used to spike-in various amounts of Covid-19 positive patient QuickExtract lysate.

Bead-LAMP using Phenol Red as colorimetric read-out (Figure S5G) was performed with WarmStart colorimetric LAMP 2x master mix (NEB) instead of the HNB containing RT-LAMP mix.

### Assessment of bead enrichment

For evaluation of the recovery rate after bead enrichment different dilutions of Twist synthetic SARS-CoV-2 RNA standard were made in HeLa cell lysate. 40 *μ*l of sample was adjusted to 100 *μ*l with QuickExtract diluted 1:1 with HBSS. Bead enrichment was performed as described for bead-LAMP. Nucleic acids were eluted with 20 *μ*l nuclease-free water for 5 minutes at 63°C. SARS-CoV-2 RNA concentrations were determined in the input (before enrichment) and eluate (after bead enrichment) by RT-qPCR.

### HomeDip-LAMP

Reactions for HomeDip-LAMP were set up as for HNB colorimetric LAMP, with final reaction volumes (excluding sample volume) being 25 *μ*l. Filter paper dipsticks (dimensions: 2×10 mm) were cut from filter paper (Fisher Scientific, cat. number 09-790-14D). Using forceps, dipsticks were dipped into 2 *μ*l of patient sample for 30 seconds, allowing the liquid to be drawn entirely onto the paper. The paper strips were then washed by rapidly submerging (‘dipping’) three times into wash solution, typically 130 mM NaCl. Sample strips were then dipped three times into the PCR tubes containing 25 *μ*l of pre-distributed HNB RT-LAMP reaction mixes. The RT-LAMP reaction was performed for 35 minutes in a water bath that was temperature-controlled by a sous-vide heater (Allpax) set to 63°C. PCR tubes were kept upright and submerged during incubation by floating pipette tip racks.

### Preparation of crRNAs for Cas12 detection

LbaCas12a guide RNAs were ordered as reverse complementary Ultramers from IDT. A T7-3G minimal promoter sequence was added for T7 *in vitro* transcription. 1 *μ*M Ultramer was annealed with 1 *μ*M T7-3G oligonucleotide in 1x Taq Buffer (NEB) in a final volume of 10 *μ*l by heating the reaction up to 95°C for 5 minutes, followed by slowly cooling down to 4°C with a 0.8°C/seconds ramp rate. One microliter of 1:10-diluted annealing reaction was used for T7 *in vitro* transcription using the Invitrogen MEGAScript T7 Transcription kit following the manufacturer instruction. RNA was transcribed for 16 hours at 37°C and purified using AmpureXP RNA beads following instructions described in (Kellner, Koob, Gootenberg, Abudayyeh, & Zhang, 2019).

### Cas12-detection of RT-LAMP product

RT-LAMP was set-up as described above and run at 62°C for 60 minutes. Meanwhile, 50 nM purified crRNA was mixed with 62.5 nM EnGen LbCas12 (NEB) in 1x NEB Buffer 2.1 and a final volume of 20 *μ*l. The RNP complex was then incubated for 30 minutes in a heat-block and kept on ice until use. For detection, 2 *μ*l of the RT-LAMP product and 125 nM ssDNA sensor (Invitrogen, DNaseAlert HEX fluorophor) were added to 20 *μ*l of the Cas12-RNP complex on ice. Reporter cleavage was monitored in real-time using a BioRad QFX qPCR cycler with measurements taken every 5 minutes for a total of 60 minutes.

### Expression and purification of HIV-1 RT

Recombinant heterodimeric HIV-1 RT (strain BH10, GenBank accession number AH002345) was expressed and purified using a modified version of plasmid p66RTB, as previously described (Boretto et al., 2001; Matamoros et al., 2005). HIV-1 RT p66 subunits carrying a His_6_ tag at their C-terminus were co-expressed with the HIV-1 protease using the *Escherichia coli* XL1 Blue strain. The resulting p66/p51 heterodimers were purified to homogeneity by ionic exchange on cellulose phosphate P11 (Whatman), followed by affinity chromatography on Ni^2+^–nitriloacetic–agarose (ProBond^™^ resin, Invitrogen). HIV-1 RT-containing fractions were pooled and dialyzed against 50 mM Tris-HCl pH 7.0 buffer, containing 25 mM NaCl, 1 mM EDTA, 10% (w/v) glycerol, and 1 mM DTT. After dialysis, enzymes were concentrated by centrifugation in Centriprep® 30K and Amicon® Ultra-4 Ultracel®-10K devices (Merck Millipore Ltd). Purity of enzymes was assessed by SDS–polyacrylamide gel electrophoresis. RT concentrations were determined spectrophotometrically by assuming a molar extinction coefficient of 2.6×10^5^ M^−1^ cm^−1^ at 280 nm. RT active-site titration was carried out as previously described (Kati, Johnson, Jerva, & Anderson, 1992; Menéndez-Arias, 1998).

### Assessment of SARS-CoV-2 detection rates (related to Discussion)

Measured RT-qPCR Cq values from clinical samples presented in Figure 1E and 1I were transformed to copies per reaction using Cq 30 = 1000 copies/reaction (determined in Figure S4B) as reference. For calculations, entire swab volumes were set to 3 ml, from which 100 *μ*l were used for RNA extractions (eluted in 50 *μ*l; 5 *μ*l of RNA per RT-qPCR) and bead-LAMP. QuickExtract lysates were prepared with 2x buffer, and 2 *μ*l were used for RT-qPCR or RT-LAMP. Copies per reaction were then transformed to equivalent copies per original sample volume used for reactions (20 *μ*l for extracted RNA, 1 *μ*l of QuickExtract crude lysate), and projected to 3 ml total swab volumes. Detection rates were calculated by dividing the number of detected samples for each procedure by the total number of detected individuals measured by RT-qPCR on extracted RNA (most sensitive method). A robust detection limit of 100 copies/reaction was used for RT-LAMP, and 5 copies/reaction for RT-qPCR. Depending on the respective purification strategy, up to 100x fold enrichment can be achieved (bead-LAMP) from 100 *μ*l original sample.

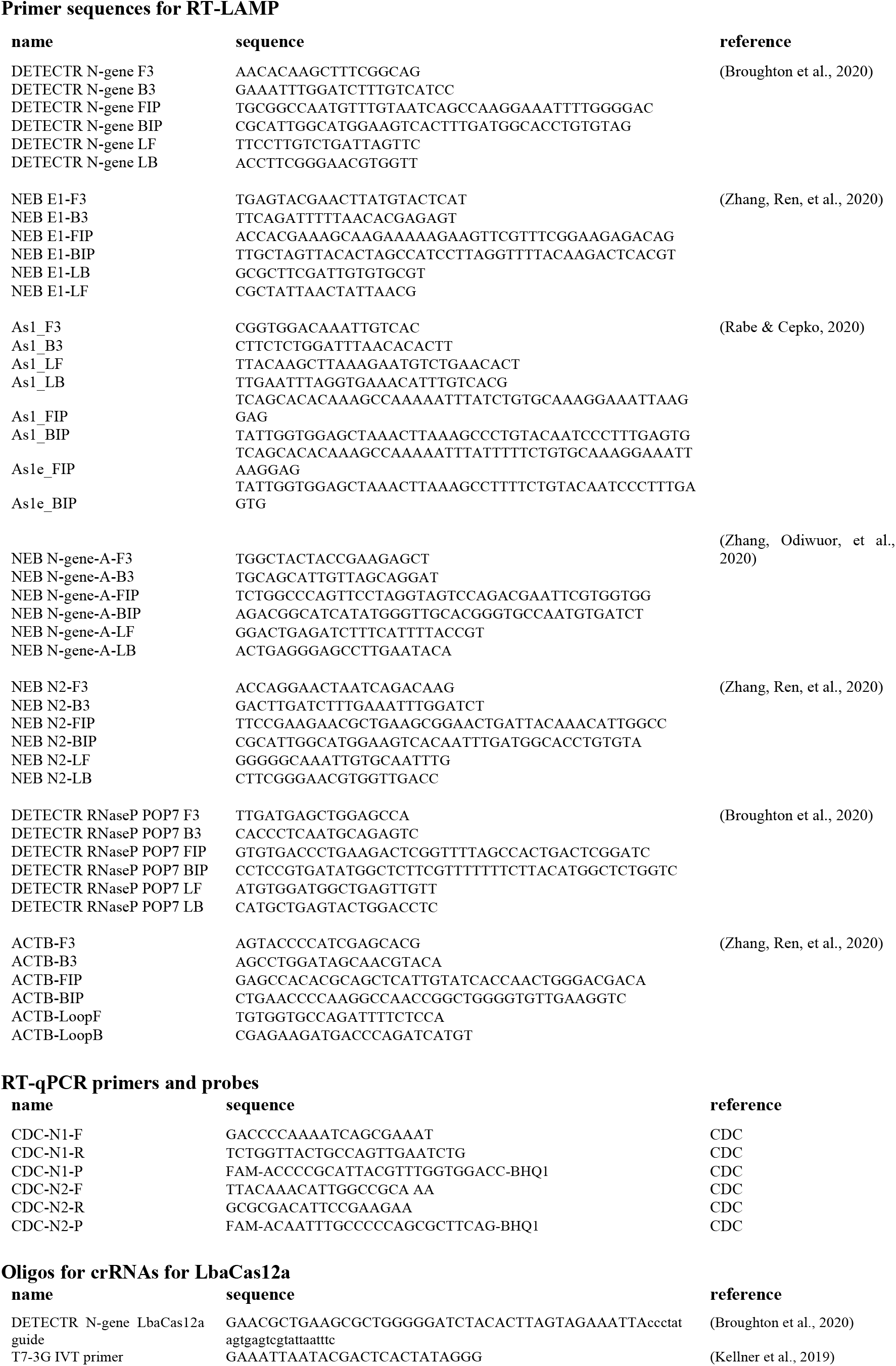

## Supplemental Figures

**Figure S1:**
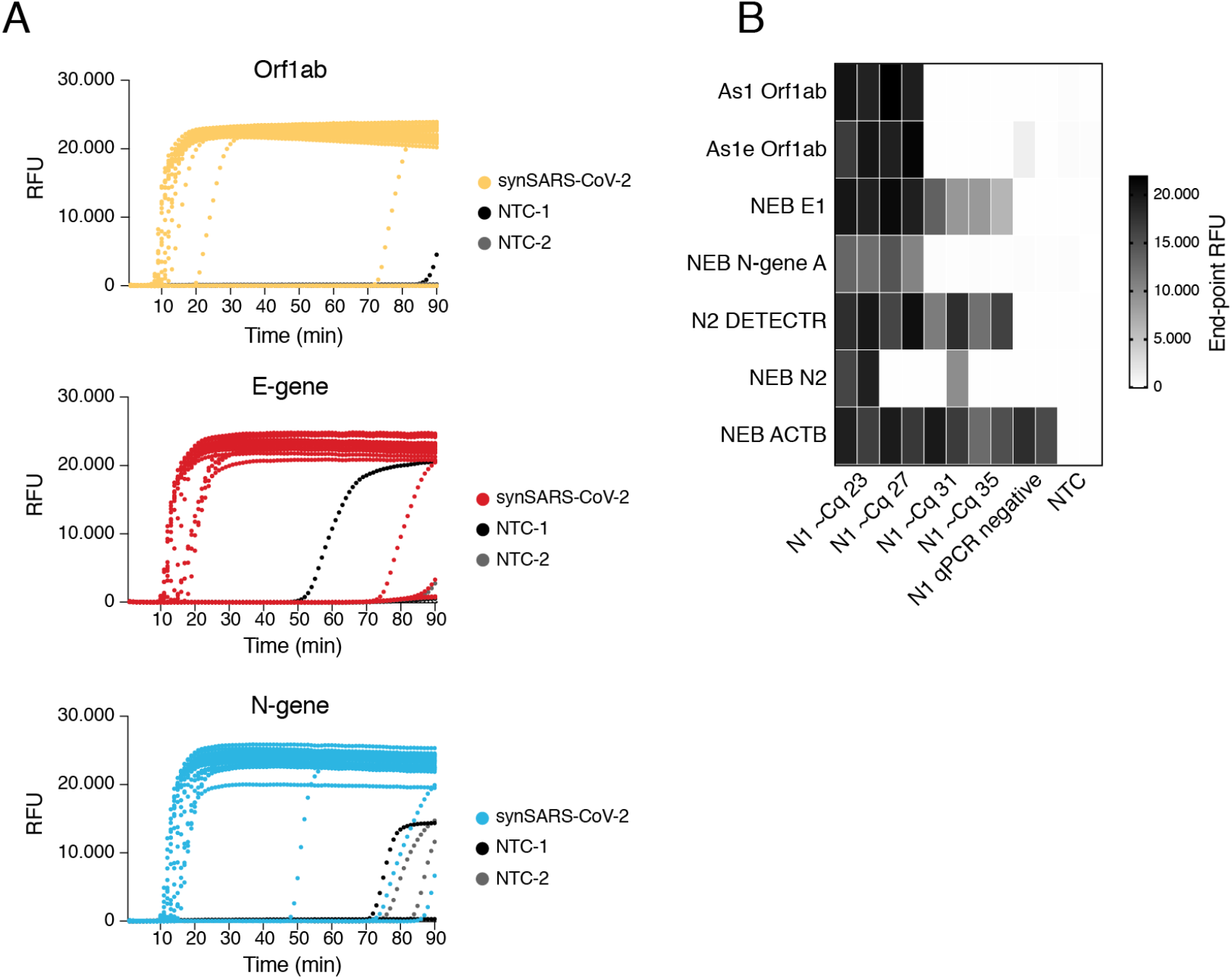
Primer performance for the detection of SARS-CoV-2 by RT-LAMP. **A)** Amplification curves (real-time fluorescent measurements; in duplicates) from RT-LAMP reactions shown in Figure 1C. Curves using synthetic SARS-CoV-2 RNA standard dilutions as input are in color (color-code indicates different primer sets; yellow: As1 Orf1ab; red: E-gene E1 NEB; blue: N-gene N2 DETECTR). Curves using non-targeting controls (NTC) as input are shown in black. **B)** Heatmap showing end-point relative fluorescence values (after 35 minutes) of RT-LAMP reactions (in duplicates; respective primers indicated to the left) using Covid-19 patient samples with indicated Cq values (determined via RT-qPCR and the N1-CDC amplicon) as input. Reactions with primers targeting the human ACTB gene served as sample quality control. All reactions were performed in duplicates.

**Figure S2:**
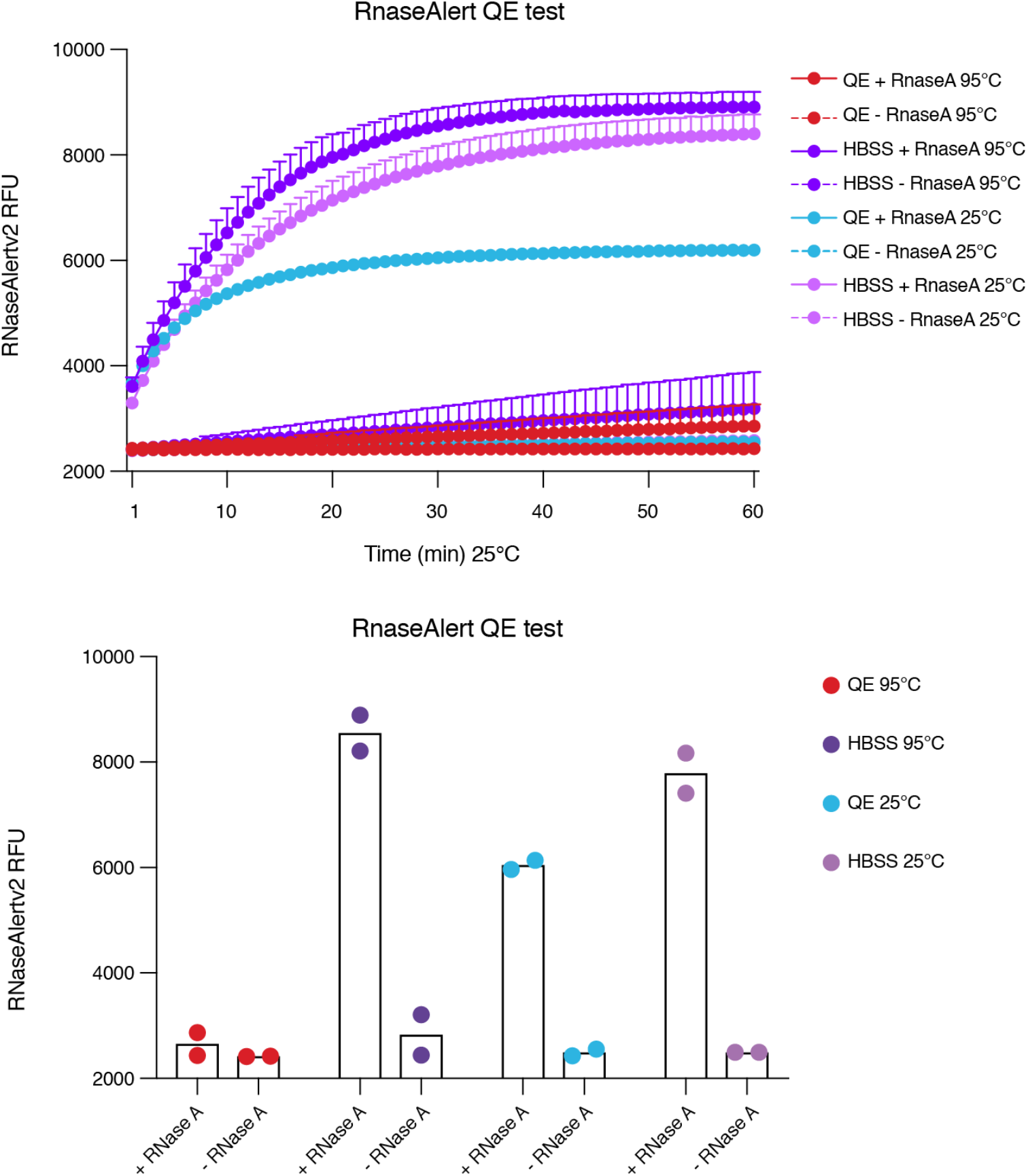
QuickExtract buffer combined with heat inhibits RNase activity. Shown are relative fluorescence values over time (upper graph) and end-point fluorescence values (lower graph) of RNaseAlert reactions in HBSS buffer or 1x QuickExtract, in the presence or absence of RNase A and with or without incubation at 95°C. All reactions were performed in duplicates.

**Figure S3:**
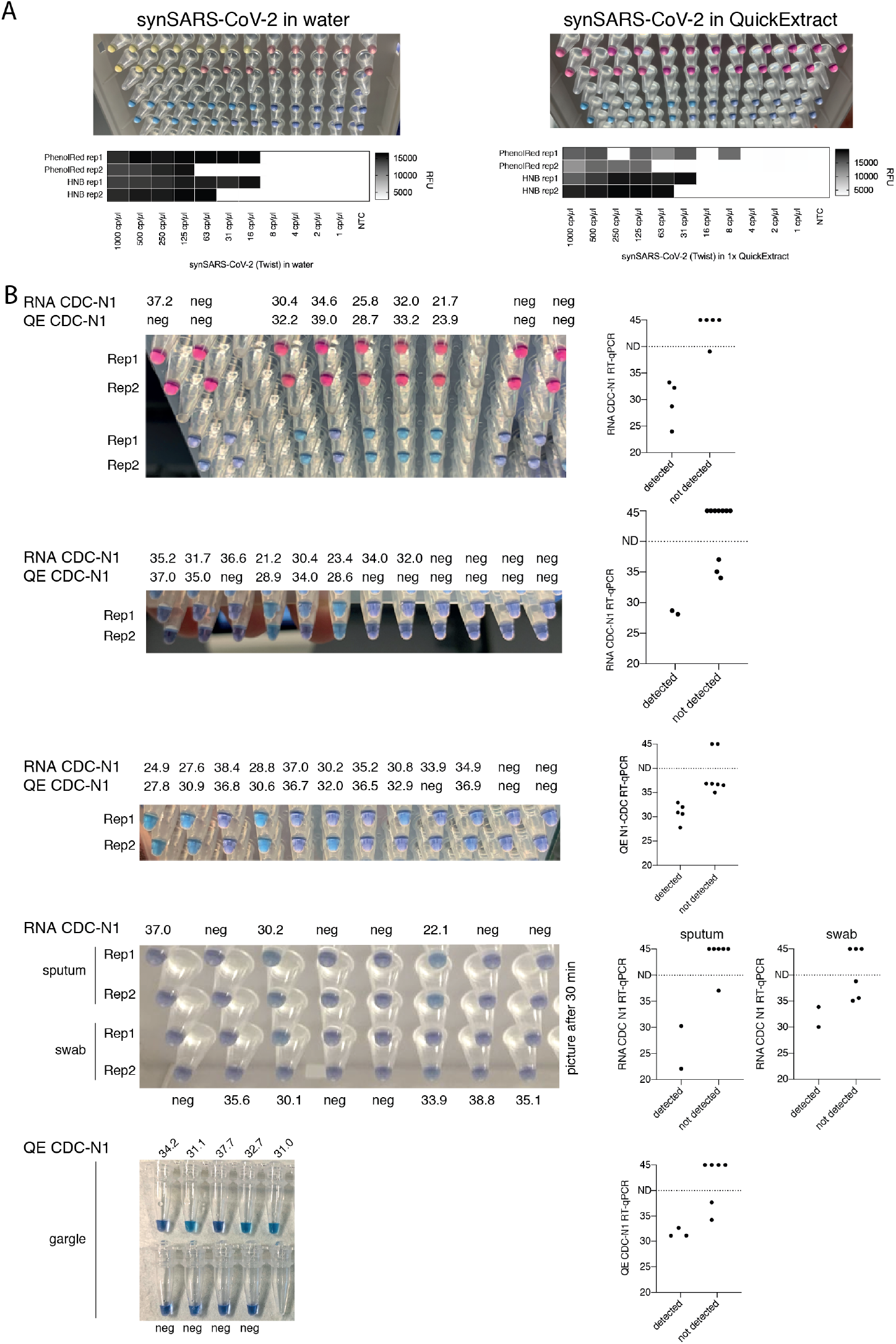
HNB RT-LAMP shows robust performance on QuickExtract-treated patient sample material. **A)** Comparison between Phenol Red and HNB colorimetric readout of RT-LAMP reactions (in duplicates) on serially diluted synthetic SARS-CoV-2 RNA in water (left) or 1x QuickExtract (right). End-point fluorescence values measured in parallel are shown in heatmaps below. While fluorescent detection indicates successful LAMP in both sample matrices, Phenol Red but not HNB colorimetric readout is inconclusive in QuickExtract buffer (right panel, top rows). **B)** HNB RT-LAMP performance across a wide range of Covid-19 patients and sample types. Images showing the HNB end-point outcome of RT-LAMP reactions on multiple Covid-19 patient samples (gargle, swab or sputum; samples indicate swabs if not otherwise stated). The respective Cq values of the individual samples (CDC-N1; QuickExtract RT-qPCR or extracted RNA RT-qPCR as indicated) are shown above or below each sample. Colorimetric RT-LAMP using Phenol Red is shown for one sample set (first from top), again with inconclusive outcome. All reactions were performed in duplicates. The HNB color-reaction was read-out at 35 minutes unless indicated otherwise. Summary dotplots for every sample set are shown to the right; samples were classified as detected or not detected based on RT-LAMP outcome and plotted against their respective RT-qPCR determined Cq values.

**Figure S4:**
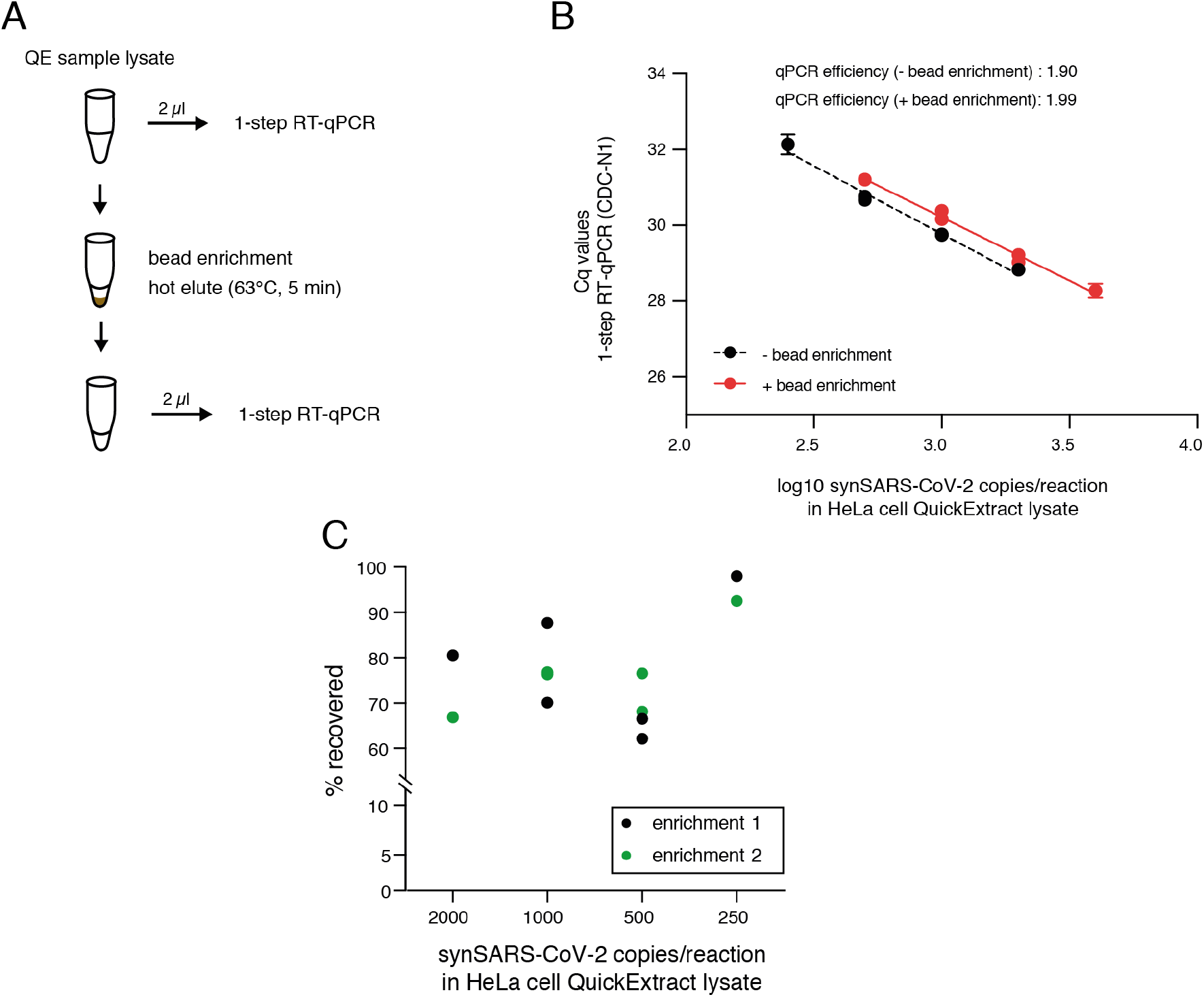
Assessment of the bead enrichment procedure. **A)** Schematic depicting the workflow to asses bead recovery performance. Synthetic SARS-CoV-2 standard was diluted in HBSS:QuickExtract lysis buffer (1:1). 40 μl were subjected to magnetic bead enrichment, followed by elution of nucleic acids in 20 μl of RNase-free water by incubation at 63°C for 5 min. 2 μl of the input (before enrichment) and the eluate (after bead enrichment) were analysed by 1-step RT-qPCR. **B)** RT-qPCR Cq values of different dilutions of synthetic SARS-CoV-2 standard before (black) and after bead enrichment (red). Duplicate experiments are shown. qPCR efficiency was calculated as 10^(−1/slope of the linear regression of datapoints). **C)** Calculated recovery (in %) after bead enrichment for the dilution series of synthetic SARS-CoV-2 RNA standard measured in B) per enrichment reaction. Replicate experiments are shown in black and green.

**Figure S5:**
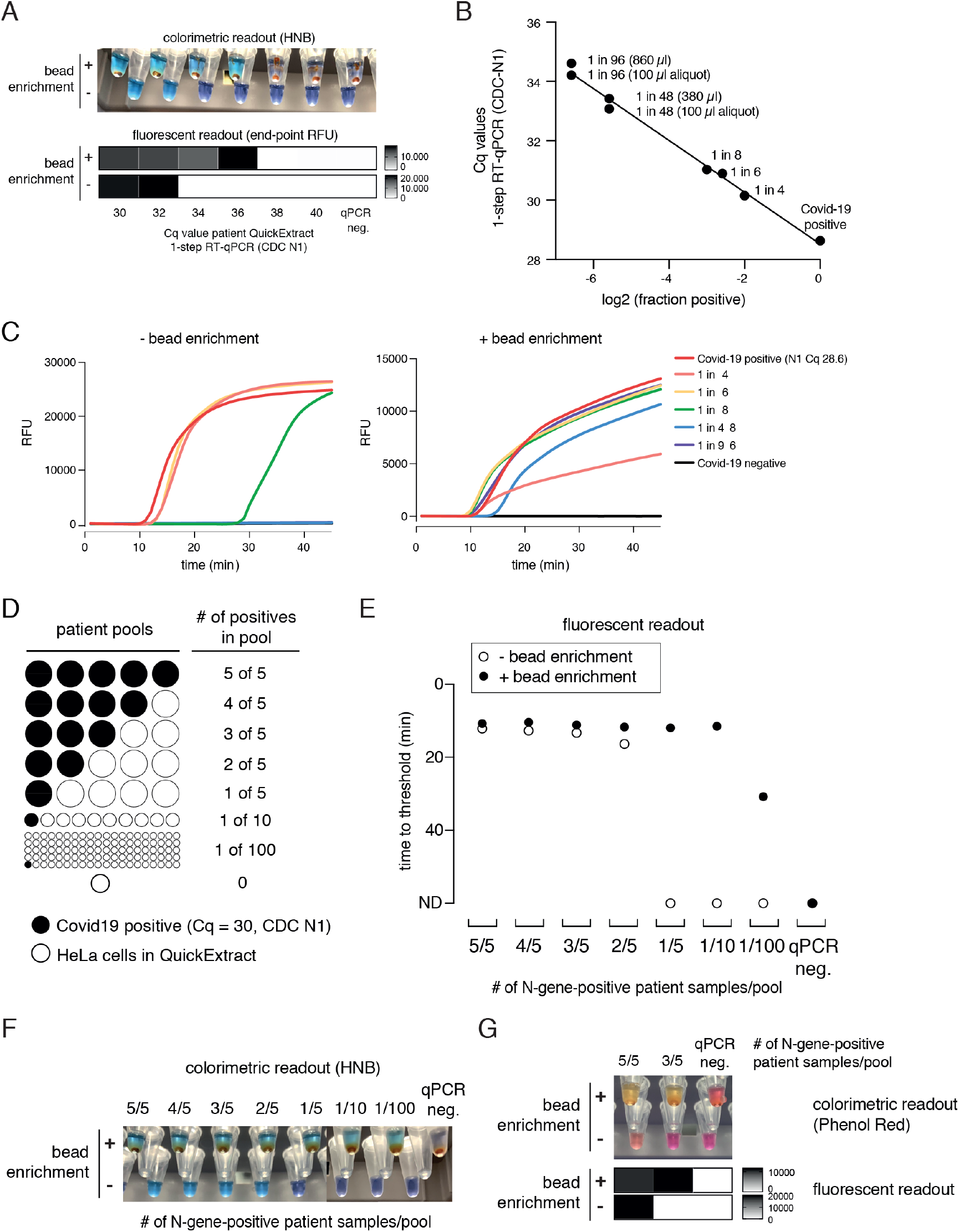
Pooled Covid-19 testing strategy using bead-LAMP. **A)** Performance of bead-LAMP on crude patient samples. The image (top) shows HNB end-point colorimetric readout and the heatmap (bottom) shows co-measured end-point relative fluorescence units (RFUs) of RT-LAMP on serially diluted patient samples in QuickExtract-prepared HeLa cell lysate, with or without prior bead enrichment. Cq values are estimated based on RT-qPCR measurement of the Cq value of the undiluted parental Covid-19 patient sample prior to bead enrichment. All reactions were performed in duplicates. **B)** RT-qPCR Cq values (CDC-N1) of gargle sample pools used in Fig 4H-J with the indicated fraction of Covid-19 positive gargle sample per pool. For the two large sample pools (pool of 1 in 96 and pool of 1 in 48), the 100 μl aliquot used for subsequent RT-LAMP and bead-LAMP was measured in addition. **C)** Readout of a real-time fluorescence RT-LAMP reaction of sample pools with indicated fraction of positive lysate without (left) and with (right) bead enrichment. RFU: relative fluorescent units. **D)** Schematic illustrating the pooled testing strategy. Eight pools mimicking different total patient sample numbers and different ratios of Covid-19-positive patient samples (0-100%) were generated from one Covid-19-positive QuickExtract patient sample (N1 RT-qPCR with Cq ~30) mixed at the indicated ratios with QuickExtract HeLa cell lysate at 20 cells/μl. **E)** Shown is the performance (measured as end-point relative fluorescence units (RFU)) of bead-LAMP (filled circles) compared to regular RT-LAMP (open circles) on the patient pools defined in D. ND = not detected within 60 minutes of RT-LAMP incubation. **F)** Images showing the endpoint HNB colorimetric readout of samples measured in E with or without prior bead enrichment. **G)** Bead-enrichment makes crude QuickExtract samples compatible with the pH-sensitive Phenol Red based colorimetric readout of RT-LAMP. Images showing the endpoint Phenol Red colorimetric readout (top) and the fluorescent readout (bottom) of two Covid-19 positive pools and one Covid-19 negative pool (qPCR negative) defined in D with (+) or without (-) prior bead enrichment.

**Figure S6:**
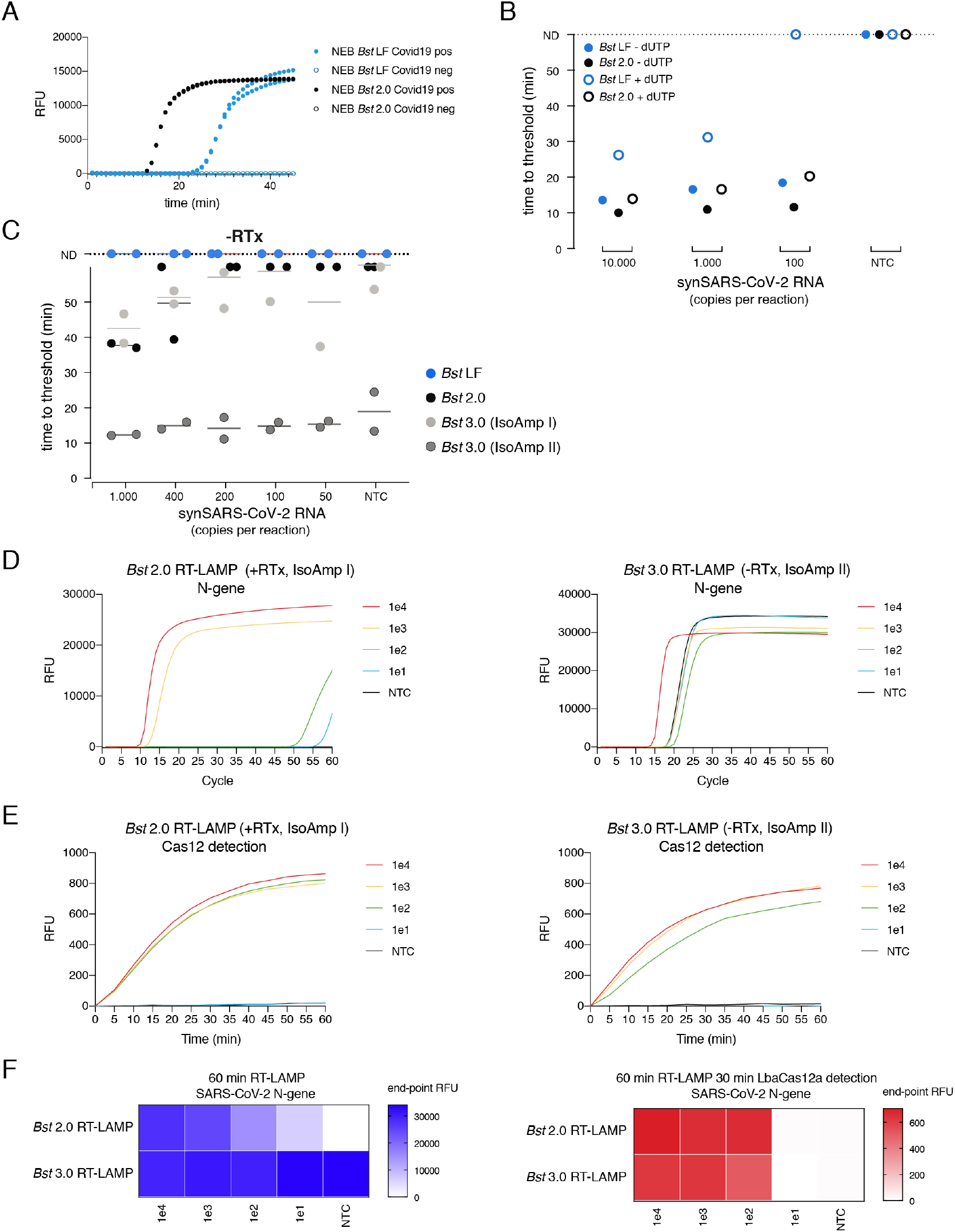
Comparison of different *Bst* polymerases for RT-LAMP. **A)** Performance of *Bst* LF (blue curves) or *Bst* 2.0 (black curves) on crude Covid-19 patient sample (prepared in QuickExtract). Amplification curves indicate real-time fluorescence measurements of RT-LAMP reactions (E1 primer set; in duplicates) using SARS-CoV-2 positive (filled circles) or SARS-CoV-2 negative (open circles) patient samples as input. **B)** Comparison of the ability of wildtype (*Bst* LF, blue) and engineered *Bst* polymerase (*Bst* 2.0, black) to incorporate dUTP during RT-LAMP on synthetic SARS-CoV-2 RNA standard. Reactions were either run under standard RT-LAMP conditions (-dUTP, filled circles), or supplemented with 0.7 mM dUTP, 0.7 mM dTTP and 1.4 mM of each dATP, dCTP, dGTP (open circles). Plotted is the ‘time to threshold’ as a measure of performance. **C)** LAMP performance (given as time to threshold in minutes) of indicated *Bst* DNA polymerase variants in the absence of a dedicated reverse transcriptase (-RTx) using diluted synthetic SARS-CoV-2 RNA (copies per reaction indicated) as template (related to Figure 6A). **D)** RT-LAMP real-time fluorescence measurements using RTx and *Bst* 2.0 in IsoAmp buffer I (left) versus *Bst* 3.0 alone in IsoAmp buffer II (right). N2 DETECTR was used as primer set for amplifying synthetic SARS-CoV-2 RNA standard (copy number per reaction is indicated; no target control (NTC): water). **E)** Shown is the collateral cleavage activity (measured as real-time fluorescent signal) by Cas12, with a crRNA targeting the N2 LAMP amplicon, upon addition of 2 μl of LAMP reactions from D) to 20 μl of Cas12 cleavage mix. **F)** (Left) End-point fluorescence values (after 60 minutes) of RT-LAMP reactions from D). (Right) Cas12-based detection of LAMP products from D) is indicated.

**Figure S7:**
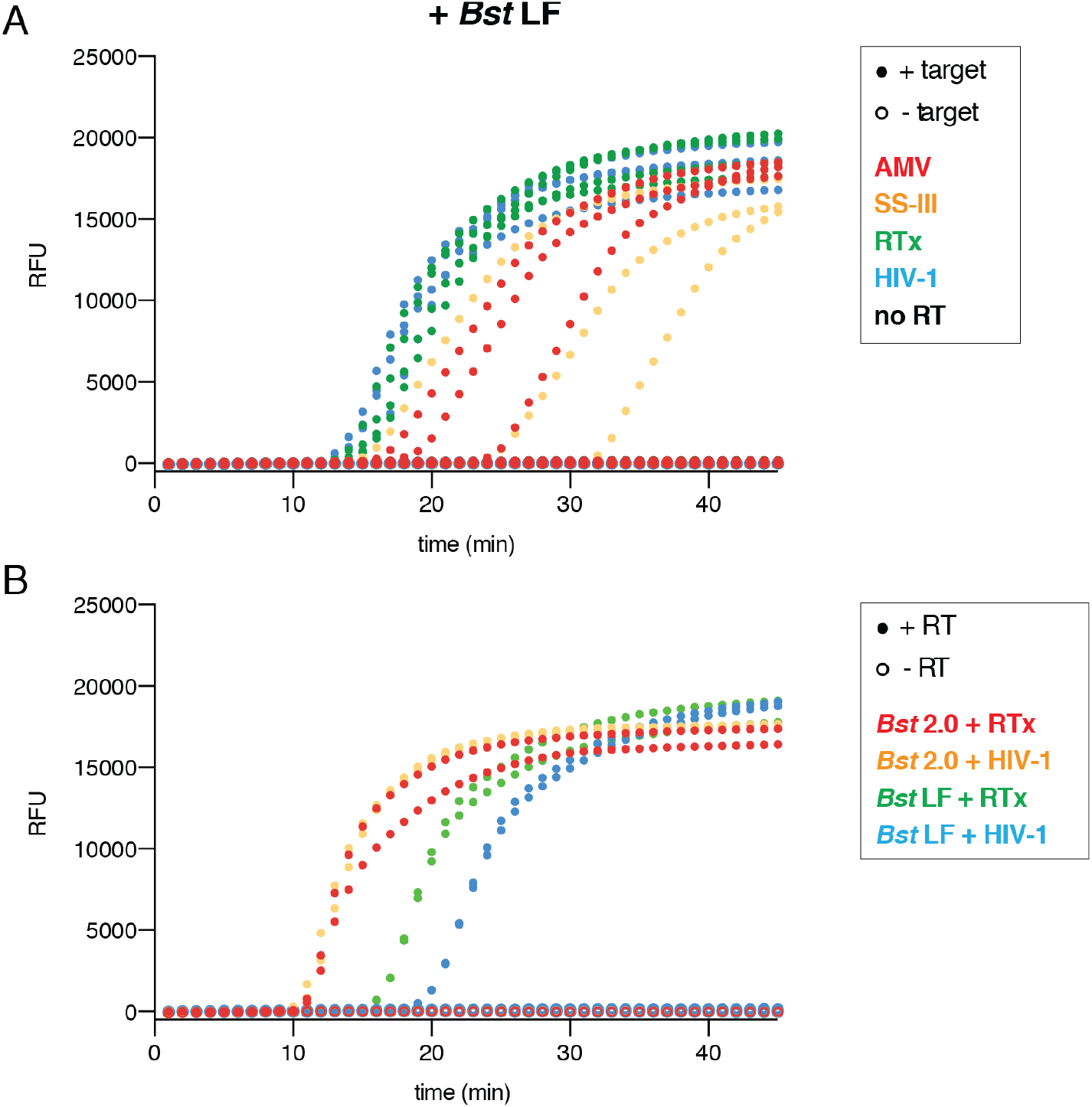
Comparison of different reverse transcriptases and *Bst* polymerases for RT-LAMP. **A)** Amplification curves (real-time fluorescent measurements; in triplicates) from RT-LAMP reactions shown in Figure 6B. **B)** Amplification curves (real-time fluorescent measurements; in duplicates) from RT-LAMP reactions shown in Figure 6D.

**Figure S8:**
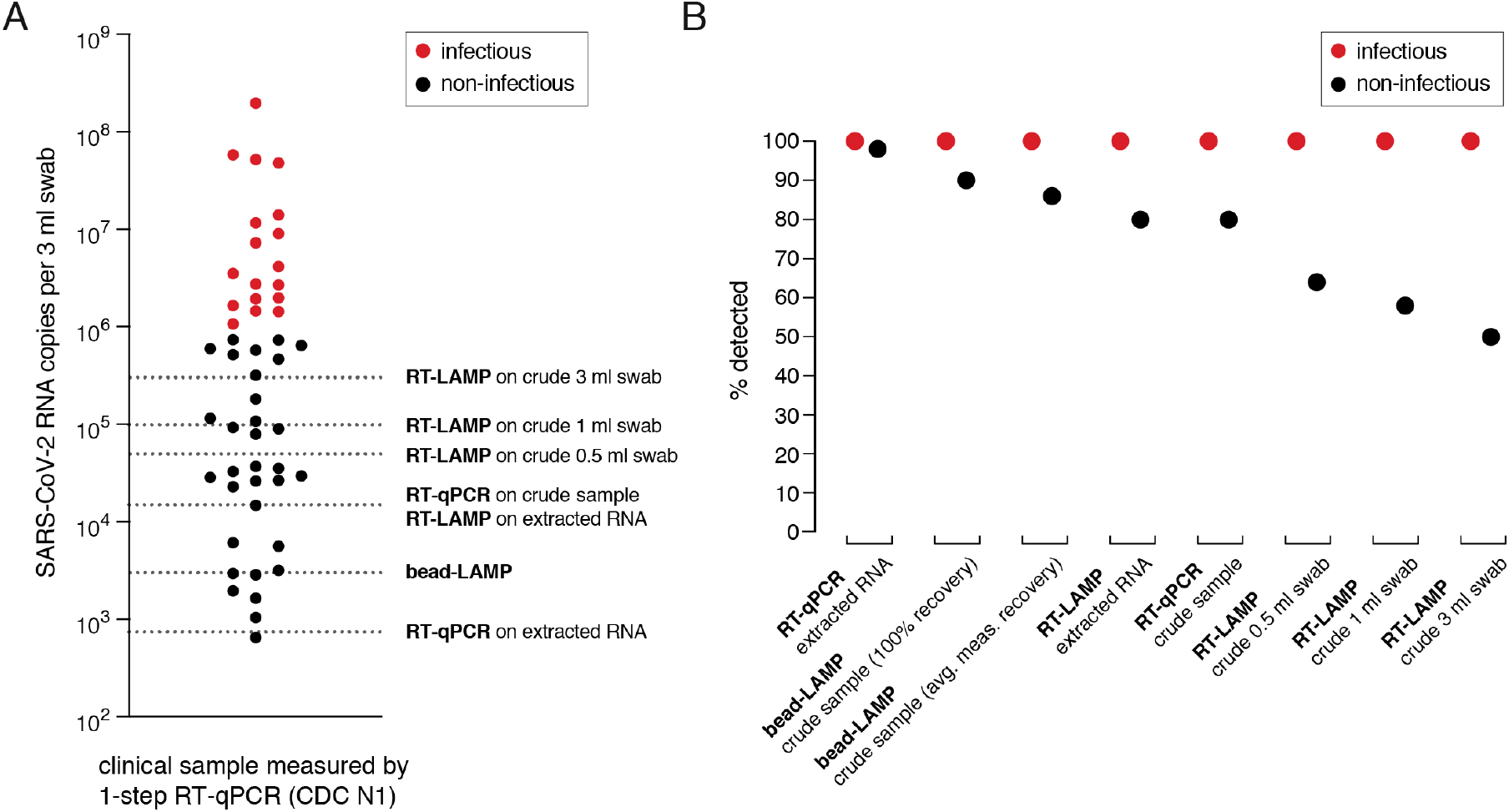
Potential of RT-LAMP-based assays for clinical SARS-CoV-2 diagnostics. **A)** Hypothetical performance of RT-LAMP-based assays based on copy numbers of SARS-CoV-2 RNA (measured by RT-qPCR) in patient samples derived from nasopharyngeal swabs in 3 ml transport medium. Copy numbers per 3 ml swab were calculated from estimated copies per reaction measured by 1-step RT-qPCR using Cq 30 = 1000 copies/ reaction as reference (Figure S4 process control). Dashed lines indicate threshold detection levels for each procedure when considering 20 μl of original sample for RT-qPCR and RT-LAMP on extracted RNA (from 100 μl original sample eluted in 50 μl), 100 μl of original sample for bead-LAMP, and 1 μl of original sample for RT-LAMP and RT-qPCR on crude lysate. Patients were classified as infectious (red) if copies per swab were >1e6 (Wölfel et al., 2020). **B)** Hypothetical detection rates of RT-qPCR and various RT-LAMP-based assays with regards to infectivity based on the data shown in A). For bead-LAMP, detection rates considering maximal nucleic acid recovery rates (100%) and measured average nucleic acid recovery rates (77%) are shown.

